# Viral rebound kinetics following single and combination immunotherapy for HIV/SIV

**DOI:** 10.1101/700401

**Authors:** Mélanie Prague, Jeffrey M Gerold, Irene Balelli, Chloé Pasin, Jonathan Z Li, Dan H Barouch, James B Whitney, Alison L Hill

## Abstract

HIV infection can be treated but not cured with antiretroviral therapy, motivating the development of new therapies that instead target host immune responses. Three such immunotherapies were recently tested in non-human primates – a TLR7-agonist, therapeutic vaccine, and broadly-neutralizing antibody – and cured a subset of animals by preventing or controlling viral rebound after antiretrovirals were stopped. However, their mechanism of action remains unknown; for example, whether they reduced the pool of latently-infected cells versus boosted antiviral immunity, and whether they acted independently or synergistically. Here we conduct a detailed analysis of the kinetics of viral rebound after immunotherapy, and use mathematical models combined with rigorous statistical fitting to quantify the impact of these interventions on viral dynamics. We find that the vaccine reduced reactivation of latent virus by 4-fold, and boosted the avidity of antiviral immune responses by 17-fold when alone and 210-fold when combined with the TLR7-agonist. In the context of later initiation of antiretroviral therapy only, the TLR7-agonist reduced latent reservoir reactivation by 8-fold, but also slightly increased target cell availability (1.5-fold). The antibody boosted immune response avidity (8-fold) and displayed no detectable synergy with the TLR7-agonist. To predict the impact of these immunotherapies in clinical trials, we calibrated a model of HIV rebound to human treatment interruption trials and simulated the effect of adding each therapy. Overall, our results provide a framework for understanding the relative contributions of different mechanisms of preventing viral rebound and highlight the multifaceted roles of TLR7-agonists for HIV/SIV cure.

## Introduction

Worldwide, over 39 million people are currently infected with HIV, and 2 million individuals are newly infected each year [1]. While combination antiretroviral therapy (ART) can suppress viral replication, preventing both transmission and progression to AIDS, it cannot completely clear the infection. A latent reservoir of integrated virus exists in long-lived lymphocytes and can re-initiate the infection (“rebound”) whenever treatment is stopped [2,3]. Consequently, current therapy must be taken for life, and new research efforts are underway to find a permanent cure for HIV [4].

Two general approaches are being taken to prevent HIV rebound and hence allow therapy to be completely stopped (“cure”). One approach, often called a “sterilizing cure”, aims to purge all remaining latent virus from the body [5], ideally re-capitulating the effects of case-studies involving bone marrow transplants (e.g. [6, 7]) or extremely early treatment initiation (e.g. [8, 9]). Another approach, often called a “functional cure”, is to instead equip the immune system with the ability to control virus that reactivates from latency, perhaps mimicking what occurs in so-called elite controllers [10] or post-treatment controllers [11]. Because of the difficulties in detecting latent virus and the lack of known immune correlates of HIV control, all current potentially-curative interventions must be evaluated by conducting treatment interruption studies, in which recipients eventually stop all therapy in a controlled manner and are monitored closely for viral rebound [12].

Here we analyze data from three recent studies [13–15] that are part of a larger effort to use therapies that stimulate the immune response (known as “immunotherapy”) to induce viral control either by clearing latent virus, boosting antiviral immune responses, or both. One component of the investigation therapy is a small-molecule agonist of the Toll-like receptor 7 (TLR7), part of the innate immune system involved in antiviral defense [16]. Another component is a “therapeutic vaccine” (administered *during* infection, as opposed to traditional *preventative* vaccines). The vaccine contains HIV (or SIV) DNA encoded in a viral vector (either Ad26, an adenovirus vector, or MVA, a modified vaccinia virus vector) and is administered in a prime-boost regimen (one vector, then the other) [17, 18] (Figure 1). The third component is the PGT121 monoclonal antibody, which targets the V3 loop of the HIV envelope protein, neutralizes a large portion of global HIV strains, and has the highest neutralizing potency of any antibody isolated to date [19, 20]. Innate immune stimulation has shown promising results in animal models and early-stage trials as a strategy to treat chronic viral infections [21, 22], supplement vaccination [23–25], or enhance the effect of monoclonal antibodies [26].

**Figure 1:**
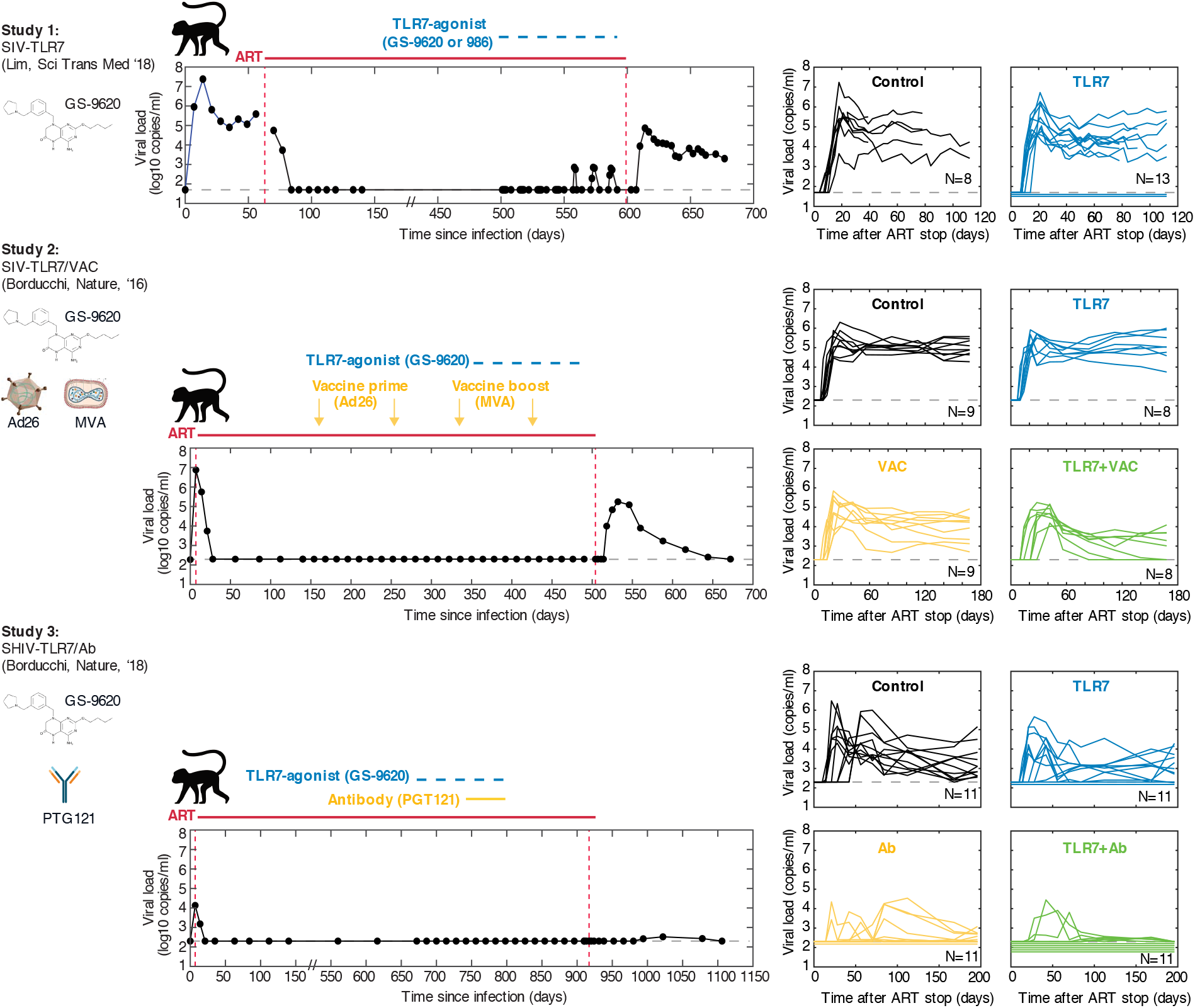
Overview of study designs for TLR7-agonist therapy with and without therapeutic vaccination or a monoclonal antibody. In **Study 1** [14], rhesus macaques were infected with SIV, and treated with ART after 9 weeks. During ART, one group of animals was administered repeated doses of a TLR7-agonist compound. ART was stopped after between 1.5-2.2 years. In **Study 2** [13], rhesus macaques were infected with SIV and treated with ART after 1 week. During ART, animals were divided into four groups, receiving a TLR7-agonist, an Ad26/MVA therapeutic vaccine in a prime-boost regimen, both, or neither. ART was stopped after 1.4 years. In **Study 3** [15], rhesus macaques were infected with SHIV (chimeric SIV with HIV envelope) and treated with ART after 1 week. During ART, animals were divided into four groups, receiving a TLR7-agonist, the PGT121 monoclonal antibody, both, or neither. ART was stopped after 2.5 years. Figures on the left show the full time-course of viral load for one example animal from each study (animal IDs: Study 1 - 156-08, Study 2 - 5888, Study 3 - 6377). Figures on the right show viral loads after ART cessation for all animals in each study. Viral load values for animals with no detectable virus are shown below the detection limit for visualization purposes only. More details of the experimental design is provided in the **SI Methods** and in the original study manuscripts.

Two of the pre-clinical studies we examine were conducted in SIV-infected rhesus macaques, a well-validated animal model of HIV infection which recapitulates HIV pathogenesis and ART response [27]. For studies involving the monoclonal antibody, animals were instead infected with SHIV, a chimeric virus consisting of the HIV envelope gene in an SIV backbone. One or two of the immunotherapies was given to animals during long-term ART, and viral levels were monitored once all therapies were discontinued. The kinetics of viral rebound were altered in many treated animals, and a subset of animals showed unprecedented responses - some animals never rebounded and appeared to have achieved a sterilizing cure, and another subset rebounded temporarily but then achieved complete suppression of virus (apparent functional cure) (Figure 1). The goal of this study was to characterize in detail the changes in rebound kinetics affected by each component of the immunotherapy. Furthermore, we wanted to compare hypotheses about the biological mechanism of the therapies, evaluate their synergy, and make informed predictions about how they may perform in human trials.

Mechanistic mathematical models are a well-established tool for characterizing and quantifying the dynamics of HIV infection within individual hosts (reviewed in [28–30]). These “viral dynamics” models are generally represented in the form of non-linear ordinary differential equations, and have been instrumental in understanding many important aspects of infection. Insights from models include the cause of viral load decline post-peak [31, 32], the lifespan of infected cells [33, 34], the rate of seeding of the latent reservoir [35], the effects of treatment with antiretroviral therapy [36–39], and investigational immunotherapies such as antibodies [40] and IL-7 [41]. Viral dynamics models can be used to statistically fit time-series data over the course of infection (especially viral load values) and to infer the values of one or more of the biological quantities represented by the model parameters. Traditionally, such fitting was done on an individual-by-individual basis using least-squares minimization methods (e.g. [40, 42]). However, individual-level fitting methods often suffer from identifiability problems when data is sparsely sampled or variables are unobserved. Different values of the model parameters may generate similar probability distributions of the observable variables, making it diffucult to compare dynamics between treatment groups. More recently, inference methods based on non-linear mixed-effects models have been developed to jointly infer parameters from groups of individuals in a population approach [43, 44]. By assuming some underlying structure to the distribution of individual-level parameters across a population, these approaches improve identifiability, and provide a formal method for testing for differences between treatment groups (e.g. [39, 45]). Parameter estimation for population-level fitting approaches can be done using either maximum likelihood or Bayesian approaches (e.g. [37, 46]).

In this paper we describe the use of mathematical modeling to understand the effects of single and combination immunotherapies on viral rebound kinetics following ART-cessation. First, we develop an augmented model of HIV/SIV dynamics which includes latent infection and an adaptive immune response. Then we analyze the dynamics of this model and investigate the theoretical identifiability of its parameters from longitudinal viral load data. We present a non-linear mixed effects statistical inference framework to estimate model parameters from the data and use this to evaluate the most likely mechanism of action of each component of the treatment.

## Results

### Development of a viral dynamics model for rebound and control

Animals who received either ART alone or augmented with the TLR7-agonist, therapeutic vaccine, monoclonal antibody, or combination immunotherapies exhibited a wide range of viral rebound trajectories (Figure 1). The standard mathematical model of HIV viral dynamics, which describes interactions only between virus and target CD4 T cells, cannot explain prominent features of these kinetics, such as a large difference between peak and set-point viral load or eventual control of infection [42]. We hypothesized that these kinetics were influenced by the induction of an adaptive immune response before and during rebound. This idea is supported by the obsevation that immunotherapy led to perturbations in interferons and interferon-stimulated genes, activation of multiple lymphocyte subsets, and expansion of cellular immune responses to viral peptides [13–15].

To infer the mechanisms of immunotherapy action across the full range of observed viral rebound kinetics, we augmented the standard model of viral dynamics to account for an adaptive immune response. In addition to modeling uninfected and infected target cells and virus, we included population of effector immune cells which suppress infection and a longer-lived precursor population which produces effectors and provides immunological memory. Our model is general enough to represent either cellular or humoral responses. We also modeled the reactivation of latently infected cells, which provides the initial source for rebounding virus. The model was specifically developed to be flexible enough to capture rebound kinetics both in the regime where latent cells reactivate frequently and rebound occurs rapidly, and in the regime where reactivation from latency is rare and there are stochastic delays until the first fated-to-establish lineage exits the reservoir [46–48]. Figure 2 shows the model schematic and Eq. (S1) gives the mathematical description. This augmented model is able to qualitatively reproduce the diverse rebound trajectories seen in data from the two studies, including rebound followed by a high set-point and rebound followed by immune control (Figure 3, S4). The **SI Methods** section details the study designs, therapies, data collection, model structure, and fitting methods.

**Figure 2:**
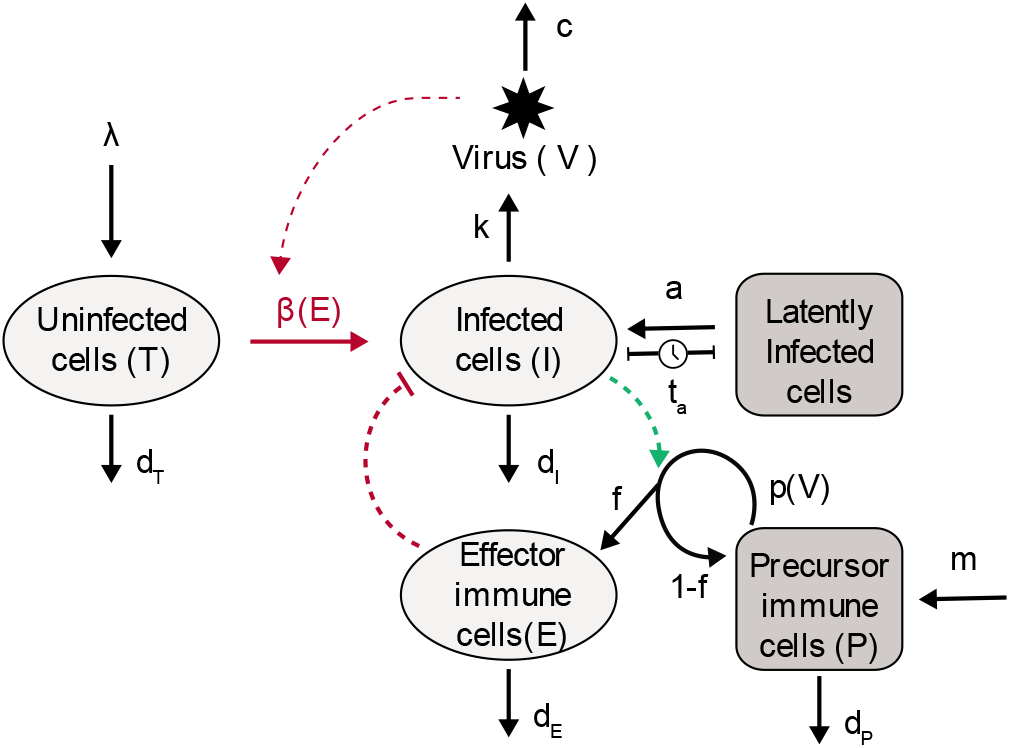
Schematic of the viral dynamics model with latent infection and an antigen-dependent immune response. Briefly, free viruses *V* enter target cells *T* (with infection rate *β*), producing infected cells *I*. Infected cells in turn release free virus (rate *k*). Long-lived precursor immune cells P which encounter viral antigen proliferate (*p*(*V*) = *pV*/(*V* + *N_P_*)) and produce short-lived effector immune cells *E*. Effector immune cells eliminate some infected cells before they contribute to ongoing infection by producing new virus (*β*(*E*) = *β*/(1 + *E*/*N_E_*)). A fraction *f* of expanded precursor cells revert to the precursor state after encountering antigen, forming immunological memory. Both uninfected target cells and precursor immune cells are produced at a constant rate (*λ* and *m* respectively). While during acute infection *m* likely represents activation of naive cells, during viral rebound, it may be dominated by reactivation of memory cells. Latently infected cells reactivate with rate *a* (or equivalently, every *t_a_* days on average) to become productively infected cells. Virus is cleared at a rate *c* and each cell type *i* dies with death rate *d_i_*. Model equations are given in Eq. (S1) and parameters listed in Table S1. More details are provided in the **SI Methods**.

### Simulation and identifiability analysis of viral rebound kinetics

The goal of this study was to understand the effects of immunotherapy by estimating the parameters of the model from the observed rebound trajectories and then comparing these parameters between treatment groups. To justify this approach, we first sought to determine which model parameters could be identified from viral load time-series alone, and to understand the expected influence of these parameters on rebound kinetics. A comprehensive identifiability analysis, detailed in the **SI Methods**, found that the combination of reservoir reactivation rate *a*, target cell replenishment rate λ, and viral burst size *k* can be adjusted to produce equivalent observed trajectories (since we only have longitudinal measures of viral load *V* and not infected cells *I*). Likewise, we can only identify the ratio of precursor immune cell production rate *m* and half-maximal inhibitory concentration of effector cells *N_E_* (since we don’t have measures of effector immune cell levels *E*). Accordingly, we fixed the viral burst size *k* and the effector half-max *N_E_* so the remaining parameters were identifiable. Furthermore, viral clearance rate, lifespans of each cell population, and the fraction of expanded immune cells that revert to memory can be estimated from other data sources, and so we fixed these parameters at experimentally-determined values (Table S1 and **SI Methods**).

The resulting model has six remaining parameters to be estimated: target cell replenishment rate λ, infection rate *β*, the latent cell reactivation rate *a*, precursor immune cell production rate *m*, maximal proliferation rate of immune cells *p*, and the half-maximally stimulating level of virus *N_P_*. We verified that this combination of parameters was formally identifiable (see **SI Methods**), and then simulated the model under systematic variations in each parameter to understand how each affects rebound kinetics. If the immune response is not strong enough, then the kinetics of this model reduce to those of previous viral dynamics models without immune responses (Figure 3A,B). The availability of target cells (λ) and the baseline viral infectivity (*β*) control the early viral growth rate, while the timing of rebound depends on the rate at which latent cells reactivate (a). The density of target cells that the virus can access (λ) also influences the eventual setpoint viral load.

**Figure 3:**
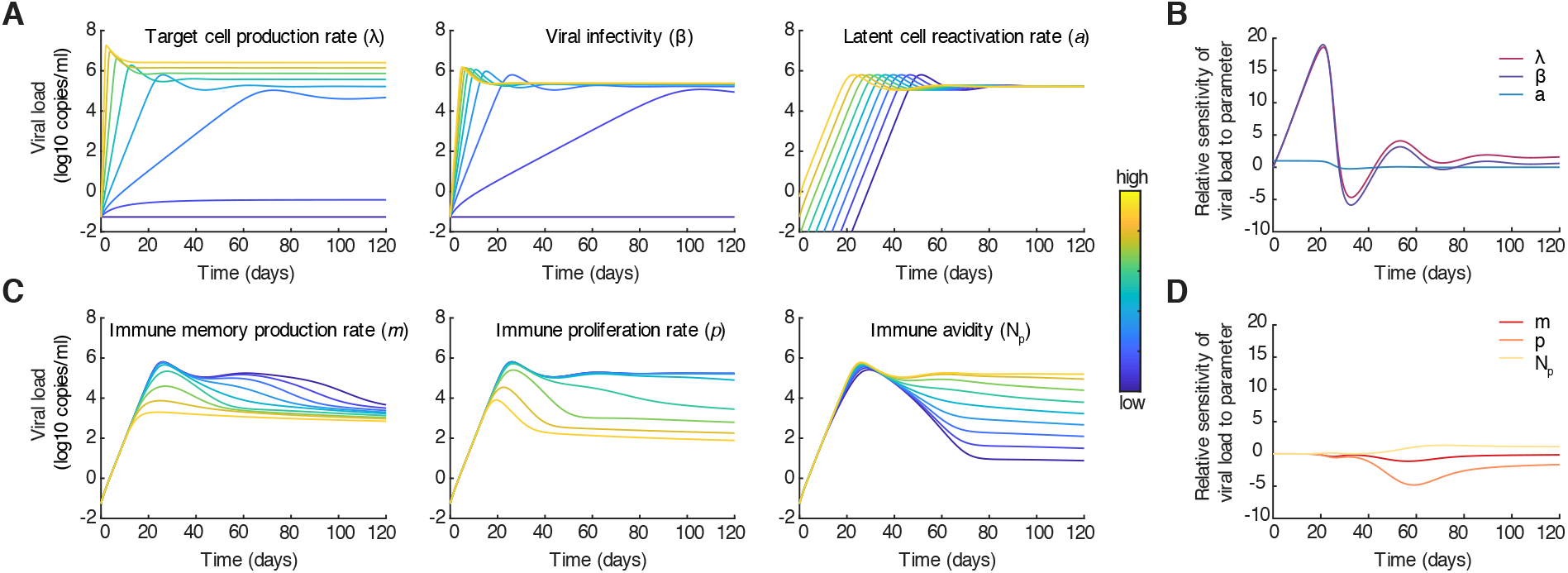
Impact of kinetic parameters on viral rebound trajectories Top row: Weak immune response. A) Viral load trajectories produced by the model for different values of either target cell production rate (λ), viral infectivity (*β*), or latent cell reactivation rate (*a*). B) Sensitivity of viral load to parameter values λ, *β*, or *a* over time. Relative sensitivity to parameter *θ* is defined as 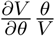. Bottom row: Strong immune response. C) Viral load trajectories produced by the model for different values of either immune memory production rate (*p*), immune proliferation rate (*p*), or immune repsonse avidity (*N_P_*). D) Sensitivity of viral load to parameter values *m, p*, or *N_P_* over time. Parameter values, when not varied and [min,max] when varied: λ = 50 [0, 500] cells mL^−1^ day^−1^, *β* = 5 × 10^−7^ [0, 20] mL copies^−1^ day^−1^, *N_E_* = 10^4^ cells mL^−1^, *d_T_* = 0.05 day^−1^, *a* = 10^−5^ [10^−12^, 10^−4^] cells mL^−1^ day^−1^, *d_I_* = 0.4 day^−1^, *k* = 5 × 10^4^ virions cells^−1^ day^−1^, *c* = 23 day^−1^, *m* = 1 [0.01, 100] cells mL^−1^ day^−1^, *f* = 0.9, *p* = 0.1 (A,B) or *p* = 1 [0.01, 10] (C,D) day^−1^, *N_P_* = 10^4^ [10^2^, 10^6^] copies mL^−1^, *d_E_* = 1 day^−1^, *d_P_* = 0.001 day^−1^. See alternate scenarios in Figure S4.

However, when viral antigen stimulates immune cells sufficiently, the immune response can curtail rebounding infection (Figure 3B,C). The level of peak viremia can be reduced by increases in the antigen-driven proliferation rate of effector immune cells (*p*), or, by increases in the rate of immune precursor cell production (*m*) (which in this model determines the size of the memory pool at the time of ART interruption). The degree of control of the setpoint viral load is determined mainly by *N_P_*, the viral load level at which antigen-stimulation is half-maximal. When *N_P_* is smaller, immune cells can still effectively proliferate even when viral load drops, thus maintaining control. The timing and rate of early viral growth - before the immune response has expanded to sufficient levels - are still controlled by λ, *β*, and *a* (Figure S4).

These results are corroborated by formal sensitivity analysis (**SI Methods**), which indicates that viral load measurements early on during rebound tend to provide the most information about the parameters related to target cells (λ), viral fitness (*β*), and reservoir reactivation (*a*), whereas measurements later on have more information about the immune response (*m, p, N_P_*) (Figure 3D).

### Estimation of immunotherapeutic treatment effects from viral rebound data

After confirming that our model can qualitatively capture a wide range of viral rebound dynamics, we used a statistically rigorous group-level fitting approach to identify the model parameters from the observed viral rebound data. Our main goal was to compare parameters between groups receiving different combinations of immunotherapy. In inference framework, baseline parameters governing the dynamics in each individual are assumed to be drawn from a shared distribution which allows for heterogeneity between individuals, known as the *random* effects. Fixed *treatment* effects alter the parameters according to the treatment each individual received. Here, the treatments we consider include the TLR7-agonist, the Ad26/MVA therapeutic vaccine, and the PGT121 monoclonal antibody. We considered animals infected with SIV and SHIV separately, as we expected that many viral dynamic parameters could differ between these virus strains. For the two SIV studies, we also included the study identity (Study 1 [14] or Study 2 [13]) as a “treatment” in order to search for systematic differences in rebound kinetics, which are most likely to be caused by the different timing of ART initiation between the studies (9 vs 1 week after infection, respectively). In addition to exploring a collection of biologically-motivated models (**SI Methods**, Tables S6 and S7), we refined our model using an iterative model selection procedure based on the Bayesian Information Criterion (BIC). Inference was performed using an implementation of the Stochastic Approximation of Expectation Maximization (SAEM) algorithm using Monolix [44, 49–51]. Instead of directly estimating *a*, the effective rate at which cells reactivate from the latent reservoir, we estimated *t_a_*, the average waiting time between reactivation events and used a formula (Eqs. (S31) to transform it into *a* to go into the model equations (Eqs. (S1)). The transformation was designed so that it worked in both the regime of frequent, deterministic reactivation associated with larger reservoir sizes and rare, stochastic reactivation from smaller reservoir sizes. The **SI Methods** section contains more details about the parameter inference and model selection.

We first examined the effects of the TRL7-agonist and therapeutic vaccine in the studies involving SIV-infected animals (Studies 1 and 2 in Figure 1). Our model-fitting procedure reliably identified effects of the immunotherapy on several model parameters, suggesting mechanisms for the efficacy of these treatments (Figures 4, S5; Table S4). We found that therapeutic vaccination reduced the rate of successful reactivations from the latent reservoir (↑ *t_a_*) by 4-fold (95% CI 2-8) and increased the sensitivity of immune cells to antigen stimulation (↓ *N_P_*) by 17-fold (95% CI 5-67). In addition, our fitting supported an interaction between the vaccine and the TLR7-agonist, resulting in an additional 12-fold increase (95% CI 3-47) in immune sensitivity to antigen (↓ *N_P_*) when both therapies were administered together, for a total 210-fold boost. We hypothesize that vaccination, alone and in concert with TLR7-agonist treatment, establishes an adaptive immune response with wide breadth which reduces the fraction of viruses archived in the reservoir which can successfully reactivate, and primes adaptive immune response to expand in the event of reactivation.

**Figure 4:**
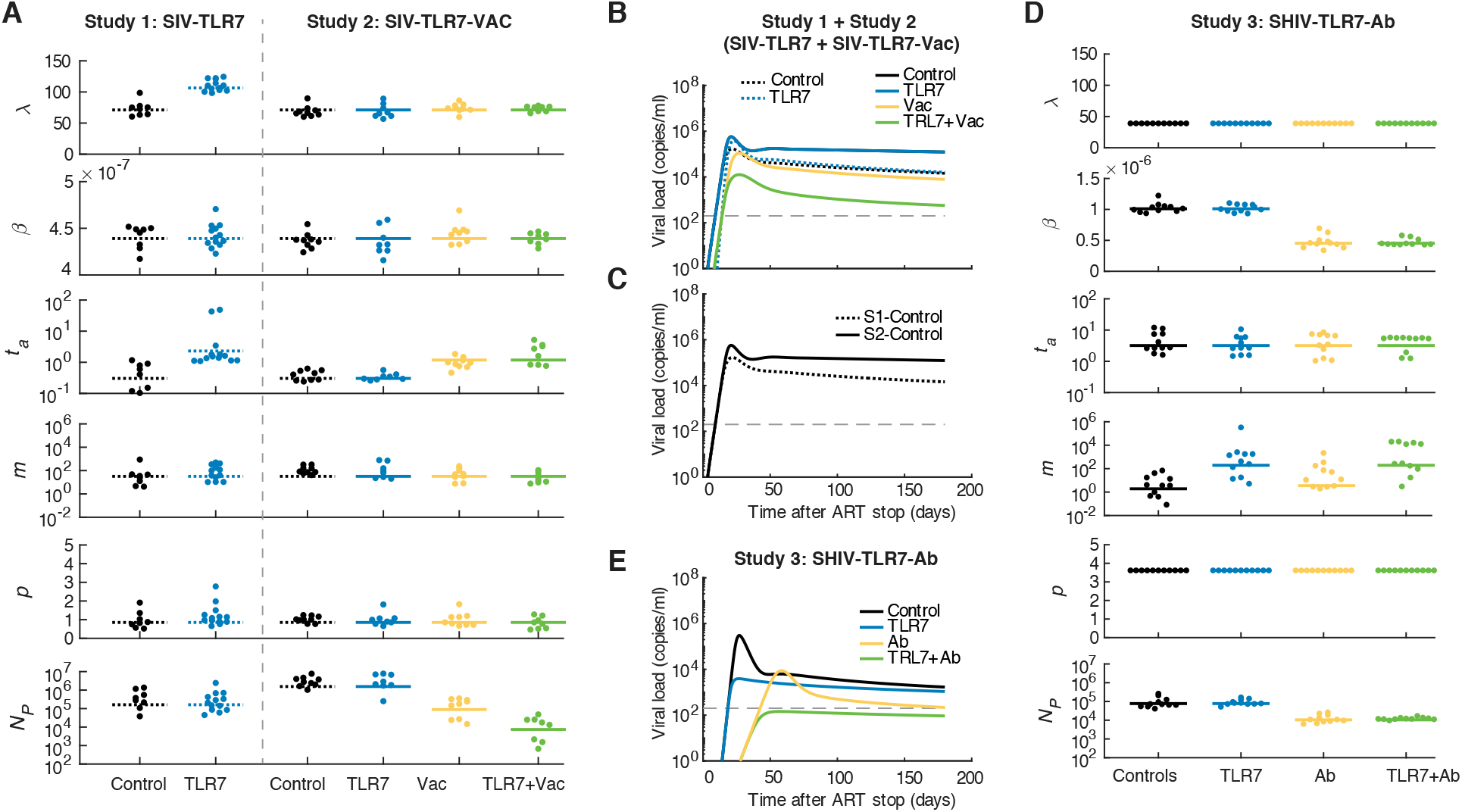
Treatment effects estimated from model fitting. A) Individual and group-level mean parameters estimated for each model parameter for the studies conducted in SIV (Study 1 and 2). B) Simulated viral load trajectories using the group-level mean value of each model parameter for all treatment groups in Studies 1 and 2. Note that since there are no group-level differences inferred between the Control and TLR7-agonist groups in Study 2, these curves are on top of each other. C) Comparing the control groups of Study 1 and 2 only. D) Individual and group-level mean parameters estimated for each model parameter for the study conducted in SHIV (Study 3). D) Simulated viral load trajectories using the group-level mean value of each model parameter for all treatment groups in Study 3. Model parameters are: λ, target cell input rate, *β*, viral infectivity, *t_a_*, time between latent cell reactivations, *m*, immune memory input rate, *p*, maximal immune proliferation rate, *N_P_*, antigen threshold for immune stimulation.

Furthermore, we identified several study-specific effects. In Study 1 [14], viral rebound kinetics supported a 10-fold elevation (95% CI 3-30) in the responsiveness of immune cells (↓ *N_P_*) in all groups, and an 8-fold reduction (95% CI 4-16) in the latent reservoir reactivation rate in the presence of TLR7-agonist treatment (8-fold increase in time between reactivations, *t_a_*). We hypothesize that the first finding is due to the longer time after initial infection that ART was started in this study compared to Study 2 (9 weeks vs 1 week), which could have allowed for the formation of a more effective memory response. Indeed, previous investigation of immunological dynamics early in acute infection indicates that the timing of ART initiation determines the strength of HIV-specific immune responses and the kinetics of the subsequent rebound [52]. The inferred reduction in the latent reservoir exit rate is consistent with the observation of large viral blips despite ART during TLR7-agonist treatment in Study 1 [14], which suggests reactivation and clearance of latently infected cells. This reduction was inferred even if we excluded the two animals who never rebounded (outliers with large values of *t_a_* in Fig. 4A).

We next examined the impact of the TLR7-agonist and the PGT121 antibody alone and in combination in SHIV-infected animals (Study 3 in Figure 1, [15]). The SHIV data exhibited greater variability than the SIV data overall: there was greater heterogeneity in the time to viral rebound and larger differences between peak and setpoint viral load. The mathematical model described the SHIV kinetics well, though the inferred residual error was larger than for SIV. We found that antibody administration produced a 2.2-fold reduction (95% CI 1.7-2.9) in viral infectivity (↓ *β*) and a 7-fold improvement (95% CI 3-16) in the responsiveness of immune cells (↓ *N_P_*) during rebound (Figure 4, S6, Table S5). While the mechanisms causing these effects remain unclear, antibody administration might eliminate more fit viral strains from the latent reservoir, leaving behind less-infectious virus. Antibody administration might also have boosted endogenous antiviral immune responses, as previously observed [53] and modeled [54].

To evaluate the robustness of our results, we used three different algorithms for selecting the optimal combination of treatment effects and for each of these we tested optimization based on both Bayesian Information Criteria (BIC) and log-likelihood (see **SI Methods**). We also varied the initial conditions for all parameters, to ensure that the algorithm did not converge on suboptimal estimates. In all cases, the models we report are robust to these variations. Despite the theoretical identifiability of all parameters we fit for, there was sometimes evidence of mutual information shared between the viral infectivity *β* and the target cell density λ. We therefore tested that an alternate model structure to our best-fit selection, which swapped the location of a treatment effect between *β* and λ, was indeed worse. The same procedure was conducted for treatment effects on *N_P_, p*, and *m*, which all describe some aspect of the immune response and hence could be hard to separate. We again found our selected model was optimal (Tables S6, S7). Before beginning the selection procedure, we defined a set of models based on biologically-motivated hypotheses about the potential effects of these treatments, and we later tested that none of these models were better than the selected one. All these results are reported in Tables S6 and S7. In Study 1 and Study 2 there was some variation in the doses of the TLR7-agonist given between animals (Table S8), and after including dose as a covariate in the model we could find no discernible influence on any kinetic parameters.

### Predicting the effects of immunotherapeutic treatment in humans

Finally, we used these results to predict how TLR7-based immunotherapies would alter rebound kinetics in human trials. Our approach was to first develop a calibrated model of HIV rebound, and then simulate the model after adding in the treatment effects (for the antibody, vaccine, etc) that we identified in the SIV and SHIV studies. To characterize HIV rebound, we assembled data from a series of clinical trials that included treatment interruptions after long-term suppressive antiretroviral therapy initiated during chronic infection, totaling 69 individuals sampled at least weekly [55]. We fit our mathematical model to these viral rebound trajectories to determine the population-level distribution of the model parameters (see **SI Methods**, Figure S7). Comparing the rebound kinetics between SIV, SHIV, and HIV (Table S9), we found large differences in the parameters estimated for different viruses. The rate of reactivation from the latent reservoir was estimated to be the highest for HIV (smallest time between reactivations, *t_a_*), followed by SIV and then SHIV. This parameter is not scale-invariant and instead depends on the absolute number of latently infected cells, so the larger body size of humans as compared to macaques likely explains the apparent difference number of cells reactivating per day. The target cell density was inferred to be the largest for HIV (↑ λ), whereas SHIV was found to have both the highest intrinsic viral infectivity (t β) and most sensitive immune response (↓ *N_P_*). The inter-individual variation in rebound trajectories (only considering control animals) was low for SIV but higher for HIV and SHIV.

To predict how a human cohort might respond to immunotherapy treatment, we next conducted simulations where we altered the baseline parameters for HIV rebound by the immunotherapy effects identified in this study. Underlying this approach is the assumption that each therapy component will have the same relative effect in humans as in macaques (e.g. a 5-fold reduction in reservoir reactivation). All single and combination immunotherapies involving the TLR7-agonist, the Ad26/MVA therapeutic vaccine, and the PGT121 monoclonal antibody were simulated in a hypothetical population of individuals, and we calculated the distribution of peak viremia, setpoint viral load, and time to rebound for each case (Figure 5, S8–S13). These simulations predict that the TLR7-agonist/vaccine combination could be effective in humans: 38% of simulated individuals rebounded above the detection threshold and then subsequently controlled viremia below it, and another 18% had no detectable viral rebound for a year after treatment interruption. The combination of TLR7-agonist/antibody treatment is predicted to result in a dramatic suppression of viral rebound in humans, primarily because the antibody’s reduction of viral infectivity (*β*) is sufficient to push the viral growth rate below the critical threshold *R*_0_ = 1 in many individuals. Supplemental figures S8, S9, S10, S11, S12, and S13 show predictions for many other hypothetical treatment scenarios in humans and macaques.

**Figure 5:**
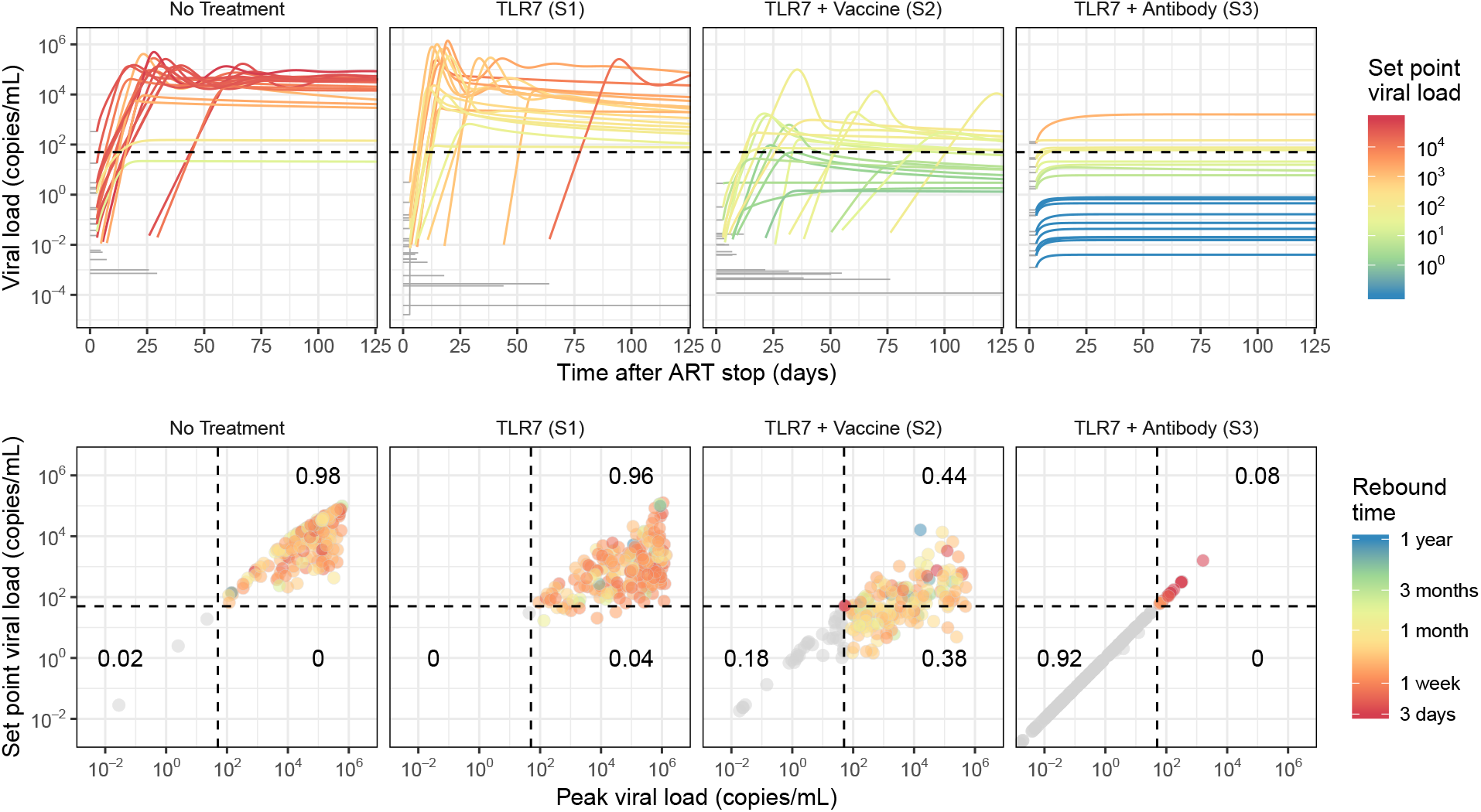
Simulated HIV rebound after immunotherapeutic treatment in humans. Viral rebound trajectories were simulated by combining baseline population heterogeneity from human HIV rebound data with treatment effects from the macaque studies. To simulate an individual rebound trajectory, a set of parameters was sampled from the human population fitting results and then the relevant treatment effects were applied. The treatment effects applied in each panel correspond to the treatment effects identified in the indicated study; for clarity the list of effects can be found in Table S10. The top row shows 20 example rebound trajectories colored by viral load. Viral load prior to successful rebound is illustrated by a grey horizontal line at the level expected according to the simulated reservoir size. Each trajectory is colored according to its viral load at one year. The bottom row summarizes 200 simulations for each treatment according to their peak and final viral loads. Each dot represents a particular individual and is colored according to the first time at which that individual rebounded (crossed 50 copies/mL). Individuals who never crossed this threshold are shown in gray. Dotted lines indicate the detection threshold for standard viral load assays (50 copies/mL) and dark black numbers show the proportion of individuals falling in the indicated quadrant. Viral rebound trajectories were simulated for one year.

## Discussion

Immunotherapy is a promising new approach to achieve long-term remission, or even cure, of HIV infection. Currently the only way to access the efficacy of these interventions is to interrupt antiretroviral therapy (ART) and observe whether viral rebound occurs. In this study we analyzed data from a collection of trials conducted in macaques to test three new immunotherapies - a small-molecule TLR7 agonist, a therapeutic vaccine (Ad26/MVA), and a monoclonal broadly neutralizing antibody (PGT121). We developed and applied a mathematical model of viral dynamics in order to characterize rebound kinetics in detail and better understand the mechanisms of action of each therapy. Using a statistically rigorous group-level fitting framework, we were able to quantify the effect of each therapy on several immunological parameters and on the latent reservoir. Moreover, we identified synergistic effects when different immunotherapies were used in combination, for example, between the TLR7 agonist and the therapeutic vaccine. Beyond elucidating these treatment effects, our analysis provides a formal way to compare and explain differences in rebound kinetics between SIV, SHIV, and HIV, and with early vs late ART initiation. Finally, our results provide a framework to translate therapeutic effects observed in animal models to treatment outcome predictions for in human clinical trials.

Across all the human and macaque studies we examined, our viral dynamics model was able to describe the rebound kinetics in most subjects very well. The main benefits of this model are that it is a relatively minimal yet mechanistic model of adaptive immunity, and that it can describe stochastic delays until viral rebound within the context of a differential equation, allowing established model-fitting procedures to be used (e.g. Monolix). We believe our approach to modeling reactivation from latency had several advantages over simpler models upon which it was based (e.g. [13, 14, 46, 48, 56]). Firstly, our model coherently captures the relationship between reservoir size and rebound, including interpatient variability in both quantities, when reactivation is rare *and* when it is common, without making an assumption *a priori* about which regime the kinetics are in. Secondly, by integrating estimation of the reservoir reactivation rate with the whole time course of rebound, we can better account for the fact that the probability that a reactivating cell gives rise to productive infection is a function of other model parameters. Long delays to rebound have been observed in individuals with very low latent reservoirs [7–9], and we expect that as the efficacy of new reservoir-reducing therapies improves, rebound dynamics in this regime will become more common.

However, our approach has several limitations. Our model fails to account for the gradual increase in set point viral load that occurs during HIV/SIV infection, characteristic of the transition to immunodeficiency. It also only describes a single viral strain and a single immune response, and hence does not account for within-host evolution of the virus nor the diverse and evolving set of lymphocyte clones responding to infection. Any observed features of viral rebound in our study subjects which reflect these or other un-accounted-for processes could have been erroneously attributed to other model parameter or treatment effects in our fitting. We do not explicitly model immune exhaustion, and this model offers an alternative to previously proposed “bistability” explanations for post-treatment control [57]. We fit our model only to viral load values, since they were the only densely-sampled quantity in these studies. As a result, many model parameters were fixed in order to improve identifiability, and absolute values of the fit parameters are less reliable than differences observed between groups. Furthermore, this limited our ability to critically evaluate the functional form of our model of adaptive immunity and estimate absolute values for immunologic quantities. In future studies, longitudinal sampling of immunological covariates should be conducted so that more detailed dynamical models of “systems immunology” in the context of infection can be developed.

Our statistical approach to model-fitting makes several assumptions that must be kept in mind when interpreting our results. One assumption is that the model parameters are similar across individuals within a group, more specifically, that they are log-normally distributed. This assumption provides additional statistical power, but if it is severely mis-specified, the results will not be reliable. Secondly, the fitting approach assumes that a treatment effect acts to alter the parameter values of all individuals in the treatment group by a fixed (multiplicative) amount. Our finding of treatment interactions (or lack thereof) must be interpreted in the context of this underlying assumption. Finally, we adapted tools designed primarily for fitting deterministic models to the problem of describing viral rebound in the regime where at least one process - reactivation from latency - behaves stochastically (see **SI Methods**). Future work might approach fitting fully-stochastic simulations of viral rebound to data while maintaining this group structure among individuals.

Cure strategies for HIV are often categorized as either sterilizing cures - which act to clear latent virus - or functional cures - which act to inhibit viral spread. However, our results suggest that individual immunotherapy agents may actually act by both mechanisms of cure, albeit incompletely, since rebound still occured in most subjects. Moreover, we found that the inferred mechanism of action could depend on details of the study design, such as the viral strain (SIV vs SHIV) and the timing of initiation and the duration of ART. Details of these findings are discussed below.

In SIV-infected animals in Study 1, administration of the TLR7-agonist alone was inferred to alter rebound kinetics by two different mechanisms of actions. Firstly, it reduced the average rate of latent reservoir reactivation, and this was the inferred mechanism of cure in the two animals who never rebounded. This conclusion from the model is supported by additional experimental data from the original study [14]. Animals treated with the TLR7-agonist had reduced levels of integrated SIV DNA, and the non-rebounders had negative viral outgrowth assays, supporting the hypothesis that this treatment reduced the size of the latently-infected population. Detectable viral “blips” concurrent with TLR7-agonist administration also suggest that this drug has the capacity to reactivate and facilitate the clearance of a significant proportion of latently infected cells. TLR7-agonist administration was also inferred to increase the density of target cells for the virus (which acted against its therapeutic potential). HIV and SIV mainly infect activated CD4+ T cells, and while the TLR7-agonist was observed to transiently alter the proportion of activated cells in several immune subsets [14], whether it can effect a long-term change in target cells has not been directly assessed. An increase in target cell production might also reflect a more generally healthy immune system, perhaps due to an immunoprotective effect of the TLR7-agonist. Note however that due to parameter non-identifiability within the model, we cannot rule out that the TLR7-agonist instead increased the viral burst size (k), which we did not allow to vary in our model.

Interestingly, the TLR7-agonist alone was not inferred to have any effect on SIV-infected animals in Study 2. The reason for this difference is unclear. The protocols for Study 1 and 2 were nearly identical, with the main difference being the time of ART initiation (after 9 weeks in Study 1 vs 1 week in Study 2). Even among control animals, there were differences between the studies: the model inferred viral load threshold for immune stimulation was 10-fold lower in Study 1 vs 2, (↓ *N_P_*), leading to a stronger immune response. This result agrees with previous work showing that later ART start time enhances the formation of memory immune responses [52]. We hypothesize that the interaction of this more-developed immune response with the latency-reversing activity of the TLR7-agonist was necessary for its reservoir-reducing action. However, the direction of this trend is not completely obvious, since later ART initiation can also lead to a larger latent reservoir [52, 58], incomplete CD4+ T cell recovery [40], and more opportunity for viral diversification and antigenic escape [59, 60]. Surprisingly, we did not infer a difference in the effective latent reservoir reactivation rate (*t_a_*) between the control groups in Study 1 and Study 2, whereas we had hypothesized that reactivation would be more frequent in the group that was infected for longer before ART start (i.e. ↓ *t_a_* in Study 1). It is possible that the increased time for reservoir seeding when ART is delayed is offset by the enhanced immune responses that also develop in that time and limit reservoir longevity. Although we think the difference in timing of ART initiation is the most likely explanation for the differences between these studies, we cannot rule out effects due to differences in the duration of ART (1.5–2.2 yrs vs 1.4 yrs) and in the sampling frequency after ART cessation (at least biweekly sampling for 6 wks in Study 1 vs only 2 wks in Study 2), which is crucial for accurately capturing the slope and peak of initial rebound viremia.

The therapeutic vaccine, administered in Study 2, was inferred to work both by reducing the rate of latent reservoir reactivation (↑ *t_a_* 4-fold) and by leading to the presence of an immune cell population which is very sensitive to rebounding virus (↓ *N_P_* 18-fold). The creation of such an effective memory immune population is a hallmark of successful vaccination. There are two possible explanations for the effect on the latent reservoir. The latent reservoir archives a large diversity of viral sequences [61–64], and after vaccination, many of them may not have able to give rise to productive infection, since they were targeted by vaccine-induced immune responses. Alternatively, vaccination might also have led to the creation of an immune response which began eliminating latently infected cells even before the end of antiretroviral therapy. Larger decreases in SIV DNA levels during ART in the vaccine-treated groups suggest at least some contribution of the second mechanism [13]. We also inferred that there was a synergistic interaction between the TLR7-agonist and vaccination, which further enhanced the sensitivity of antiviral immune responses (extra 12-fold decrease in *N_P_*). This effect is consistent with the role of TLR7 as a bridge between innate and adaptive immunity [65].

In the SHIV study involving the TLR7-agonist and the antibody, the data are generally more challenging to interpret. Overall, the amount of variability is much larger; viral loads display more erratic patterns that the model has difficulty explaining, and we obtained a best-fit error term more than twice as large as the SIV data. The inferred mechanism of action of the TLR7-agonist was different in this context: the only effect the model identifed was an increase in the size of the memory pool of antiviral immune responses present at the time of ART cessation (100-fold increase in *m*). One possible explanation for this observation is that TLR7 activation leads to the permanent expansion of SHIV-specific immune responses just as it might have led to the expansion of the target cell population in the SIV context. Whether this expansion was non-specific or instead required antigenic-stimulation from a latency-reversing effect of the TLR7-agonist is not clear, but we did not identify any reservoir-reducing effect of this therapy. The fact that we did not identify an effect of the TLR7-agonist on target cell replenishment in this study could be explained by SIV and SHIV likely targeting slightly different cell populations for infection [66]. Due to limits of parameter identifiability, we cannot separate the inferred effect of the TLR7-agonist on the memory pool from an enhancement in the per-cell viral inhibitory potential of immune effector cells (*N_E_*). Previous systems-immunology analysis suggests that outcomes in this study are connected to NK cell function, and TLR7 signaling plays a critical role in NK cell function [15], at least via cytokines produced by dendritic cells [67].

The inferred effects of the PGT121 monoclonal antibody were surprising. We found that the antibody boosted immune responses against rebounding virus (7-fold ↓ *N_P_*), similar to the therapeutic vaccine administered in Study 2. This could have been a direct vaccinal effect of the antibody, as has been observed in other studies [53, 54, 68], although no evidence for enhanced CD8+ T cell responses was observed [15]. Alternatively, if the antibody eliminated certain viral strains from the reservoir via antibody-dependent cellular cytotoxicity [69], perhaps the only viral strains that avoided elimination and could cause rebound were mutated strains that were actually more sensitive to immune responses mounted during rebound. The antibody was also inferred to reduce the intrinsic infectivity of rebounding virus(↓ *β*), which is difficult to explain since the antibody is no longer present when ART is withdrawn. This could also be a selection effect: perhaps only strains with lower levels of viral gene expression avoided clearance during antibody administration. Our best-fitting model did not identify an effect of antibody treatment on the rate of latent reservoir reactivation, in contrast to the the mechanism of action hypothesized in the original study [15]. Roughly, this is because the timing of rebound in the animals who rebounded despite antibody administration was not significantly different than in control animals. Consequently, the parsimonious explanation for the effect of the antibody did not include reservoir reduction, and the lack of rebound in some subjects was inferred to be a functional rather than sterilizing cure. However, we cannot completely exclude an effect on the latent reservoir, as one competing model including this effect had only slightly worse statistical support (see Table S6). No synergy between the TLR7-agonist and the antibody could be detected.

Finally, by applying the immunotherapy effects inferred by our model to a calibrated model of HIV rebound, we generated hypotheses for the outcomes of these agents in human trials. This method has the benefit of accounting for intrinsic differences in rebound kinetics between SIV or SHV and HIV, but still makes the assumption that relative changes in parameters can be translated between systems. While the specific outcome predictions should be interpreted with caution, there are several important qualitative findings of these simulations. Most importantly, patients with high peak viral load may eventually control, which clearly complicates the design of human trials. In addition, we predict that therapies which move closer to viral control in humans will also lead to higher inter-patient variability in rebound dynamics, meaning large trials sizes may be needed. The simulations also suggest that in many patients, rebound may occur but give rise to dynamics that are controlled long-term below the limit of detection for the most common HIV RNA assays. Overall, if the therapeutic effects of these treatments transfer to a human context, our results suggest they will dramatically alter the course of viral rebound.

There are many opportunities for improvement in the design of future ART interruption studies in either humans or animals, which would optimize the knowledge that could be gained from similar model-based analyses. Most importantly, frequent sampling of viral load after ART cessation (ideally twice weekly), continuing post-peak and anticipating potential delays in rebound, is extremely important for model identifiability. Dynamical models could be integrated with an analysis of immune correlates of protection by expanding the collection of longitudinal immune covariates that are collected. In order to accurately relate rebound delays to changes in the reactivation rate from the latent reservoir, it is neccessary to have good estimates for drug washout kinetics, which could be quantified with a study of very short-term ART. In macaques, we recommend that all trials use stocks of genetically-barcoded virus, which would allow for more direct estimates of changes in the latent reservoir reactivation rate due to novel therapies [56, 70].

## Acknowledgments

This work was supported by NIH grants DP5OD019851 (ALH), P01AI131365 (ALH, JBW), and P01AI131385 (ALH, MP), and P30AI060354 (Harvard CFAR; ALH, JG), as well as funding from the French National Institute for Research in Computer Science and Automation (Inria) through an International Associate Team award (MP, ALH). We thank Jesse Fajnzylber, Ronald Bosch, Evgenia Aga and the Cure Transformative Science Group for assistance with accessing HIV rebound data from the AIDS Clinical Trials Group (ACTG) studies. The ACTG is supported by NIH UM1 AI068636, and their Statistical and Data Management Center by NIH UM1 AI068634. We are grateful for feedback on the analysis provided by Yves Levy, Rodolphe Thiebaut, and Martin Nowak.

## Supplementary Information

### Methods

#### Animal Data

The design of the studies we analyzed is explained in detail elsewhere [13–15], but summarized here. All studies were conducted in rhesus macaques, which were infected intrarectally with the SIVmac251 virus (Studies 1 and 2) or SHIV-SF162P3 (Study 3) and later treated with the combination antiretroviral therapy (ART) regime of tenofovir, emtricitabine, and dolutegravir (TFV/FTC/DTG). In the Study 1, all animals were given ART starting 65 days after infection, and were treated with ART for at least 500 days, before ART was stopped. During ART, some animals were additionally administered repeated doses of a TLR7-agonist (treatment group), while others received a placebo (control group). The study was divided into several phases and arms, in which the treatment groups received slightly different courses of the TLR7-agonist therapy (see Figure 1A-B, Table S3). Overall, there were 8 control animals and 13 TLR7-agonist treated animals. In the Study 2 (SIV-TLR7-Vac), all animals started ART after only 1 week of infection, and were treated with ART for ~500 days before stopping. Animals were divided into four groups of 8-9 individuals each - one control group who only received ART, one group who additionally received the TLR7-agonist (10 doses 2 weeks apart), another group who additionally received a prime-boost vaccine regimen (2 doses of Ad26 followed by 2 of MVA, each 12 weeks apart), and a fourth group who additionally received both the TLR7-agonist drug and the vaccine regimen. In Study 3, SHIV-infected animals began ART after a week of infection and were treated for ~900 days before stopping. Animals were divided into 4 groups of 11 animals each. One group received the TLR7-agonist (10 doses 2 weeks apart), another received the PGT121 monoconal antibody (5 doses 2 weeks apart, co-inciding with the last 5 doses of the TLR7-agonist), another received both the antibody and the TLR7-agonist, and there was also a control group.

In all animals in both Study 1 and Study 2, there was at least 2 weeks between the time the last TLR7-agonist dose was given and when ART was interrupted, which was chosen to insure that the drug had washed out of the system by the time ART was stopped. Similarly, in Study 3 there was 16 weeks between the last antibody dose and ART cessation. This way, any change in viral rebound kinetics caused by the intervention must be due to a permanent perturbation made to the system, and not a direct antiviral effect of the immunotherapy. In all studies, viral load was measured every 3-4 days after ART cessation. Additionally, viral load values were measured during acute infection and during ART administration. In all cases viral load values below the detection limit of assays (50 copies/mL in Study 1 or 200 copies/mL in Studies 2 and 3) are censored.

In Study 1 with the TLR7-agonist only, all but two animals experienced persistent viral rebound. Two animals in the TLR7-agonist treatment group never experienced detectable viral load after ART was stopped. These represent the first observed cases of a potential sterilizing cure in this SIV system. In Study 2 with TLR7-agonist and vaccine, all animals initially rebounded, but three animals treated with the combined immunotherapy eventually re-suppressed virus below the detection limit. For the SHIV-infected animals in Study 3, transient low-level rebound was seen in 3/11 TLR7-treated, 4/11 antibody(Ab)-treated, and 5/11 TLR7+Ab treated animals, whereas the complete absense of rebound occured in 1/11 TLR7-treated, 2/11 Ab-treated, and 5/11 TLR7+Ab treated animals. However, even in the control group, 6/11 animals experienced viral loads reduced to near or below the detection limit towards the end of the study follow-up period (~ 196 days). This level of post-treatment control was never seen in SIV-infected animals in the control groups, highlighting the need to treat animals infected with each viral strain separately. The rebound trajectories of all animals are shown in Figure 1 (C-H).

#### Model development

The basic viral dynamics model used to describe HIV infection before and during antiretroviral therapy can also be used to describe the rebound of infection when treatment is stopped (which displays similar kinetics to acute infection). This often-used model reproduces many aspects of infection kinetics, such as exponential increase in viremia after initial infection or rebound, declining viral load after a peak is reached, and eventual stabilization at a “set point”. However, it cannot describe the diversity of viral rebound trajectories seen in these studies - such as large declines in viral load from peak to setpoint or eventual post-rebound control - and does not explicitly consider viral latency nor antiviral immune responses. To address these issues, we developed an augmented model of HIV/SIV infection dynamics which incorporated ideas from multiple different existing models of various viral infections (Figure 2) [46, 71–76]. We previously used this model in a preliminary analysis of a subset of this data [13].

The model we used is described by a system of ordinary differential equations that track changes in the levels of uninfected (T) and infected (I) target cells, free virus (V), and precursor (P) and effector (E) immune responses over time (Figure 2):

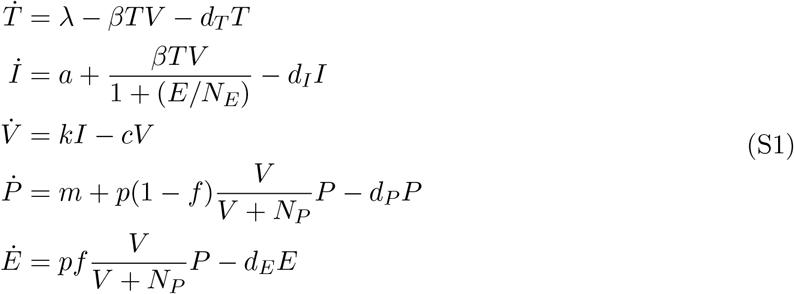

This model assumes that infection is well-mixed throughout the blood and other lymph tissue, ignoring any spatial structure or compartmentalization. All variables are expressed as concentrations (per mL of plasma). Model variables and parameters are summarized in Table S1.

Susceptible, uninfected target cells (*T*) are produced at a constant rate λ and die at a rate *d_T_*. These cells are assumed to be CD4+ T cells, but may only be a subset of the total CD4+ population. Although the specific phenotype of CD4+ T cells that confers susceptibility is not completely clear, it is known that activated cells are more susceptible to infection than resting cells, and that only a small fraction of all CD4+ T cells are productively infected even at peak viremia (more may be abortively or latently infected). Here we ignore heterogeneity in infected cell subpopulations.

New infections occur proportionally to the density of free virus (*V*), target cells (*T*), and the infectivity rate *β*. Infected cells (*I*) release virus at rate *k* and die at a rate *d_I_*. Free virus is cleared at rate *c*. We do not explicitly track latent infection, since it only significantly impacts infection levels when viral loads are very low, but instead use parameter *a* to describe the rate at which latently infected cells reactivate to produce productive infection (which is necessary to kick-start rebound). This rate incorporates the number of cells latently infected with intact virus as well as their per-capita rate of reactivation and the probability that they produce a lineage that escapes extinction and reaches detectable viremia.

**Table S1:** Variables and parameters of the viral dynamics model (Equation (S1), Figure 2). More details on parameter meanings and values is given in the **SI Methods** text. Fixed parameters that differ for humans vs macaques are given in Table S9.

| Description |  | Units |  |  |
| --- | --- | --- | --- | --- |
| **Variables:** |  |  |  |  |
| $T$ | (Uninfected) target cells | cells mL<sup>-1</sup> | | |
| $I$ | (Productively) Infected cells | cells mL<sup>-1</sup> | | |
| $V$ | Free virus | RNA copies mL<sup>-1</sup> | | |
| $P$ | Precursor immune cells | cells mL<sup>-1</sup> | | |
| $E$ | Effector immune cells | cells mL<sup>-1</sup> | | |
| **Fixed parameters:** |  |  | Value | Source |
| $d_T$ | Death rate of target cells | day<sup>-1</sup> | 0.05 | [77, 78] |
| $d_I$ | Death rate of infected cells | day<sup>-1</sup> | 0.4 | [13, 14, 52] |
| $d_P$ | Death rate of precursor immune cells | day<sup>-1</sup> | 0.001 | [77, 79–81] |
| $d_E$ | Death rate of effector immune cells | day<sup>-1</sup> | 1 | [78, 82, 83] |
| $c$ | Virus clearance rate | day<sup>-1</sup> | 23 | [84] |
| $k$ | Virus production rate | virions cells<sup>-1</sup> day<sup>-1</sup> | 50 000 | [85] |
| $f$ | Fraction of effector immune cells that don't revert to memory | 1 | 0.9 | [86] |
| $N_E$ | Effector concentration at which half-maximal inhibition occurs | cells mL<sup>-1</sup> | 10 000 | arbitrary |
| **Estimated parameters:** |  |  |  |  |
| $\lambda$ | Production rate of target cells | cells mL<sup>-1</sup> day<sup>-1</sup> | | |
| $\beta$ | Viral infectivity | mL copies<sup>-1</sup> day<sup>-1</sup> | | |
| $a$ | Latent cell reactivation rate | cells day<sup>-1</sup> | | |
| $m$ | Production rate of precursor immune cells | cells mL<sup>-1</sup> day<sup>-1</sup> | | |
| $p$ | Maximum proliferation rate of immune cells | mL copies<sup>-1</sup> day<sup>-1</sup> | | |
| $N_P$ | Viral load at which half-maximal proliferation occurs | copies mL<sup>-1</sup> | | |

We include a relatively general model of an antiviral immune response, which could represent cellular or humoral effects. Long-lived precursor immune cells (which includes both naïve and memory subsets) are produced at a baseline rate m. If this model were used during acute infection, m would be related to the frequency of naive precursors, whereas during viral rebound, m is dominated by the reactivation of memory cells formed during acute infection. In response to antigen (assumed here to be free virus, but could instead be infected cells), these cells are stimulated to proliferate at an antigen-dependent rate. The maximum proliferation rate is p and half-maximal proliferation occurs at viral load *N_P_*. The functional form of this relationship is derived from considering the chemical reaction kinetics involved in antigen binding/presentation (e.g. [87]). A fraction *f* of all proliferating cells become short-lived effectors (*E*), while the remaining fraction will return to a long-lived memory state (*P*). Long-lived immune cells die at rate *d_P_* and short-lived effectors die at rate *d_E_*. Effectors reduce the rate at which actively infected cells are produced (↑ *β*), with half-maximal inhibition occurring at an effector concentration *N_E_*. This could represent an effector population that acts to inactivate free virus, protect target cells from infection, or kill early-stage infected cells before they start producing virus. Alternatively, we could also have modeled effectors as killing virus-producing infected cells *I* (↑ *d_I_*). This is the main role assumed for CD8+ T cells, which are known to be a major component of anti-HIV immunity. However, previous work has shown that the rate of viral load decay during ART (determined by *d_I_*) is independent of CD8 levels, which is only consistent with either a non-lytic CD8 effect or the presence of an early stage of infected cells that are not yet targeted by cytolytic immune responses [75]. For simplicity, we have chosen to model non-lytic immune effects, but either model has similar qualitative behavior during rebound.

During ART, we assume infection is completely blocked (*β* = 0), and that treatment is given for long enough that virus and cells reach steady states, which we take as the initial conditions at the time of ART interruption:

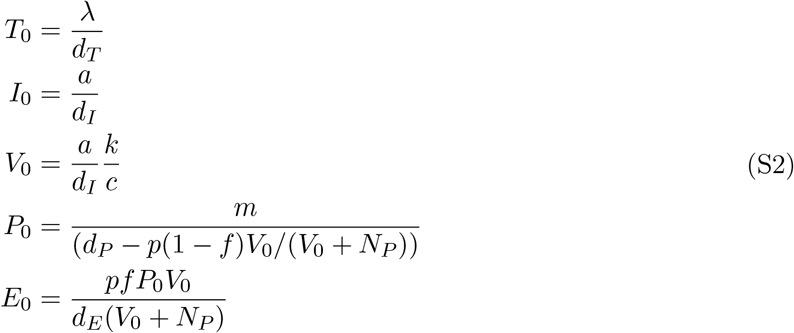

#### Model parameters and identifiability

The data available for fitting this model consists only of longitudinal values of viral load (*V*), and therefore all parameters of the model are unlikely to be identifiable. While some longitudinal values of total CD4+ T cells were collected, the relationship between this number and target cell density *T* is unclear for the reasons discussed in the previous section, and, this measurement is notoriously noisy in non-human primates sampled under anesthesia. Actively infected cells (*I*) are difficult to quantify separately from forms of latent or defective infection. Characterization of antiviral immune responses (related to *P, E*) was only done once before and after rebound.

We conducted both analytic and numeric investigation of the model to determine principled ways to reduce the number of parameters to be estimated from the data. We simulated the model under a wide range of parameter conditions, systematically varying one parameter at a time, to understand the role that each parameter played in the viral rebound trajectories (Figure 2, Figure S4). In addition, we applied the Exact Arithmetic Rank (EAR) approach implemented in Mathematica (**IdentifiabilityAnalysis** package) [88, 89] to determine the formal identifiability of the model parameters, given free virus level measurements are perfect and continuous in time.

When only viral load is observed, one of the parameters from each of the sets {λ, *a, k*} and {*N_E_, m*} is always non-identifiable. Therefore we fixed viral burst rate to *k* = 5 × 10^4^ virions/cell/day[85], and immune-response efficacy *N_E_* = 10^4^ cell/day. The latter value is chosen arbitrarily, since inhibition by immune cells is extremely difficult to measure quantitatively in experiments. However, the choice is inconsequential and only results in a scaling of the inferred *m* value (which we henceforth assume to be in arbitrary units).

While most other parameters are theoretically identifiable, many are practically impossible to infer from the available data. The fast time-scale of virus relative to infected cells makes the viral clearance rate c practically non-identifiable, and so we fixed this value to *c*=23/day based on plasma aphoresis studies [84]. Cell death rates are very hard to identify from viral loads during active infection, and didn’t have a large influence on viral load kinetics within a certain range, so we fixed values from isotope-labeling studies in the literature (/day) as 0.05 for target cells, 0.4 for infected cells, 0.001 for precursor immune cells, and 1 for effector immune cells. Target death rate was estimated based on the estimated death rate of a fast subpopulation of CD4+ memory T cells in rhesus macaques (0.05 in uninfected animals, 0.1 in those with high SIV loads) [77] and of activated memory CD4+ T cells in humans (0.08) [78]. Infected cell death rates were taken from the rate of viral load decline during ART for SIV observed in previous analyses [13, 14, 52]. The death rate of precursor immune cells, which actually represents the net decay combining cell death and homeostatic (antigen-independent) proliferation, was roughly estimated from turnover rates of slow-proliferating memory CD8+ T cells in uninfected macaques (0.0025) [77], decay of human CD8 responses to yellow fever (0.006) [79] and smallpox (0.0002) [80] vaccination, and decay of murine responses to LCMV (<0.0005) [81]. The death rate of effector immune cells was estimated from LCMV infection in mice (0.4) [82, 83] and acute mononucleosis infection in humans (0.8) [78]. We assumed the fraction of proliferating effectors that return to a long-lived memory state was (1-*f*)=0.1, consistent with experiments in multiple animals (reviewed in [86]), and results were insensitive to values other than those very near 0 or 1.

This left a model with six remaining unknown parameters: *β*, λ, *a, m,p, N_P_*. We repeated the formal identifiability analysis with this reduced model. We confirmed this reduced model is locally identifiable, and when *m* is fixed the model is globally identifiable. To understand what features of the viral rebound kinetics would be most informative of each of these parameters, and to understand how the density of samples may impact identifiability, we assessed the sensitivity of viral load (*V*) to each parameter (generically denoted *θ*) over time by evaluating 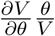, taking into account the role of the parameter in both the initial condition of the system and the subsequent time evolution (Figure 2, Figure S4).

#### Re-parameterization of model to bridge stochastic and deterministic regimes

The stochastic nature of viral reactivation means that when latent cell reactivation is rare, there can be long wait times until the first cell reactivates and thus long delays in rebound. Deterministic models of viral dynamics like Equation S1 cannot capture this wait time behavior without extraordinarily low values for the reservoir reactivation rate *a*, since they assume instantaneous input from latently infected cells (Supplemental Figure S1). In order to avoid underestimating a in individuals with delayed rebound and to correctly capture inter-individual variation in reactivation, we re-formulated our viral dynamics model to also capture reactivation in the stochastic regime. This required four changes. First, instead of directly estimating *a*, we estimate *t_a_*, the time of the first viral reactivation and interval between successive reactivations, and associate it with an a value that would have given rise to *t_a_* in expectation. Second, since the transmission of infection between cells is also an inherantly stochastic process, every cell that reactivates from the latent reservoir might not establish an exponentially-growing infection. Some cells may instead only produce low levels of infection that fluctuate before going extinct. The probability that a reactivating latent cell survives this stochastic extinction, *p*_surv_, connects the stochastic process of reactivation to the deterministic rate *a*: if reactivations occur on average every *t_a_* days, the first succesful reactivation is expected to occur after *t_a_*/*p*_surv_ days. Accounting for this establishment probability, which is a function of all other model parameters (via *R*_0_), also allows us to separate treatment effects which act directly on the reservoir reactivation rate from effects on other parameters which happen to change the establishment probability. Third, in order to convert the occurance of a single succesfully reactivated infected cell to units of concentration in the peripheral blood (which our equations track), we need a scaling factor that takes into account total blood volume (which differs between humans and macaques). This is accounted for with the initial condition during stochastic reactivation, *I*_1_. Fourth, to account for the time delay between when antiretroviral therapy is stopped and when drug concentrations decay to a low enough level that a reactivated cell could seed a growing chain of infections we allow for a washout time, *t*_wash_. The **Derivation** section details the full re-formulated model that simultaneously accounts for the stochastic and deterministic reactiation regimes and was ultimately used in the fitting. Parameters used in this reformulation are given in Table S2.

#### Model fitting

The same model fitting procedure was used for all animal studies. Studies 1 and 2 were analyzed together, but Study 3 was analyzed separately, since we assumed (and later verified) that the baseline parameters values would differ significantly between SIV and SHIV.

The model in Equation (S1) was fit to the longitudinal viral load data to estimate the values of the parameters. Briefly, in this framework, the parameter values for all individuals in the population are assumed to be drawn from a common distribution, and the goal is to estimate the mean and variance of this hyper-distribution. To ensure positivity, all parameters are estimated in log-transformed scale, denoted as 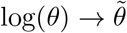. Individuals in the study may receive different treatments, and the mean parameter value may be shifted by each treatment in a different way. More explicitly, this statistical model assumes that the value of parameters *θ_i_* = (*β_j_, λ_i_, t_ai_, m_j_, p_j_, N_Pi_*) in individual *i* can be broken down into the following components

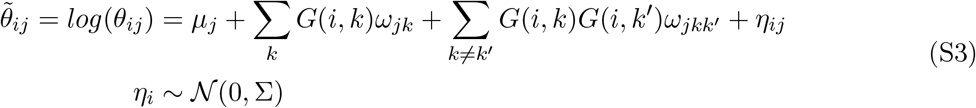

where *j* ∈ {*β, λ, t_a_, m, p, N_P_*}.

The first term *μ_j_*, known as the fixed effect, is the mean value of parameter 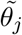 across the population in an individual who does not receive any of the treatments. The logical matrix *G*(*i, k*) describes the ‘treatments’ each individual received. If individual *i* received treatment *k*, then *G*(*i, k*) = 1 whereas it is 0 otherwise. In the analysis of Study 1 and 2 (SIV, TLR7, Vaccine), *k* = 1…3, as we consider an effect of the TLR7-agonist, of the therapeutic vaccine, and of the study identity (1 vs 2, which may influence parameters due to different initiation timing and duration of ART). In Study 3 (SHIV, TLR7, antibody), there are two treatments (*k* = 1, 2), the TLR7-agonist and the antibody. The terms *ω_jk_* describe the *treatment effects*: treatment *k* modifies the mean of parameter *j* by an amount *ω_jk_*. We also consider the interactions between these treatments that are identifiable by design, i.e. TLR7-agonist × therapeutic vaccine, TLR7-agonist × SIV study identity and TLR7-agonist × antibody. The terms *G*(*i, k*)*G*(*i, k′*)*ω_jkk′_* describe the *interaction effects*: combination of treatment *k* and treatment *k′* modifies the mean of parameter *j* by an amount *ω_jkk′_*. The final term, *η_ij_*, is known as the *random effect*, and describes the amount by which the observed parameter value *θ_ij_* differs from the expected mean in the treatment group. The *random effects* for each individual *i, η_i_*, are assumed to be normally distributed with diagonal variance-covariance matrix Σ = Diag(*ϵ_j_*)_*j*=1…5_, such that they are independent for each parameter that appears in Equation (S1).

For example, in the analysis of the SIV data, the parameter for the time between reactivation events from the latent reservoir *t_a_*, may be affected by TLR7-agonist, therapeutic vaccine administration, the study identity, and interactions between these interventions. After log-transforming 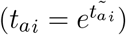, the equation for the components of this parameter becomes:

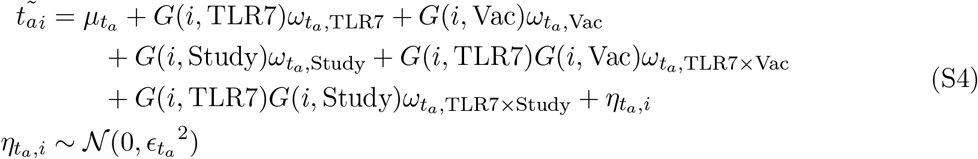

Data are always observed with measurement error, and we defined a residual error model such that the observed log_10_ viral load of patient *i* at time *t*, denoted *Y_i_*(*t*), is normally-distributed around the true viral load 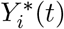 described by the model (Equation (S1)). We assume that the error has two parts: a constant error term (*ζ*_1_) and a term proportional to the log_10_ of the true viral load (*ζ*_2_) so that the observed log_10_ viral load can be written as

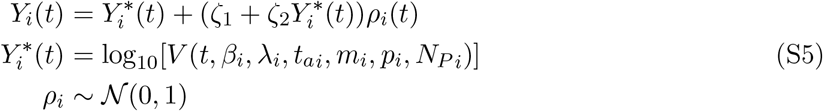

This error model is referred to as a “combined error model” and has been shown in many other studies to approximate the distribution of biomarkers measured as concentrations in blood [90]

Finally, many observed viral load values in these studies are left-censored, because they are below the detection limit of available assays (either 50 or 200 copies/mL, depending on the study). All these data points were included in our fitting, but treated formally as censored data.

We used the Monolix software version 2018R2 [44] to estimate the values of all the model parameters (*μ, ω, ϵ* and *ζ*) by maximizing the likelihood of the data given the model and parameters [44]. The software uses a frequentist version of the stochastic approximation expectation maximization (SAEM) algorithm [91]. SAEM is an iterative algorithm that essentially consists of constructing Markov chains that converge to the conditional distributions of the parameters given the data. The final parameters estimates are given by the mean parameters values over the iterations during the smoothing phase of the Markov chain, and their standard errors the uncertainty of the estimated population parameters. Standard errors are calculated via estimation of the Fisher Information Matrix [49], which is derived from the second derivative of the log-likelihood evaluated by importance sampling [92]. We did not incorporate any prior knowledge about the values of any parameters and therefore did not implement a penalized likelihood strategy. All the statistical aspects of this fitting approach has been described elsewhere [93].

#### Model Selection

In order to focus on the strongest effects and control the number of hypotheses, we determined the location of treatment effects in each dataset by employing an iterative model selection approach. Starting from a model without treatment effects but with random effects on each parameter, we applied two independent variable selection methods. First, we applied the method Stepwise Covariate Modeling (SCM) [94]. SCM has two phases: each covariate’s potential effect on each parameter is examined, and the effect leading to the best improvement in the target criteria is incorporated in the following iteration. Next, when no further improvement is possible, each previously introduced effect is removed if it fails to contribute to an improvement in the target criteria. Second, we applied the method COnditional Sampling for Stepwise Approach based on Correlation tests (COSSAC) [95]. The COSSAC method also has two phases and is similar to SCM but less exhaustive. COSSAC introduces the covariate most correlated with the random effects of a parameter into the next iteration of model fitting. When this simpler incorporation procedure stops yielding an improvement in the target criteria, a backward elimination is performed as in SCM. We used both the likelihood and Bayesian Information Criterion (BIC) as target criteria. After each selection procedure, we verified that the sign of a treatment effect was inferred unambiguously, so that it could be assigned a clear biological interpretation. To do so, we used a Wald test at level 5% to determine if treatment effects differed significantly from zero, *ω_jk_* ≠ 0.

In the SIV TLR7 ± vaccine data, we ran the selection procedure on each of the five treatment covariates: TLR7-agonist, Ad26/MVA therapeutic vaccine, study identity (Study 1 vs Study 2) and the two available interactions: TLR7-agonist × vaccine and TLR7-agonist × study identity. In the SHIV TLR7 ± antibody data, we ran the selection procedure on each of the three treatment covariates: TLR7-agonist, PGT121 monoclonal antibody, and the interaction TLR7-agonist xantibody. First, random effects were assumed on all six parameters of the model. However, in the SHIV TLR7 ± antibody data, it was not possible to achieve practical identifiability of all random effects [96], thus we decided to remove the ones on *p* and λ as they were found to be related with the non-full-rank of the Fisher Information Matrix.

Repeated fitting suggested that the SAEM algorithm employed by Monolix converged to a global maximum. We ran 100 final estimations with high number of iterations in the burn-in, exploratory, and smoothing phase of the SAEM algorithm. Each fitting produced consistent final estimates, and we selected the best in term of maximization of the log-likelihood for further interpretation. These highest-likelihood estimates of the various treatment effects were considered as final and presented in the article. We confirmed that this last model was the best in term of BIC compared to all other models tested. We also investigated over-parametrization and thus overfitting by checking the ratio between the largest and the smallest eigenvalue of the Fisher Information Matrix, which remained small, in both cases this ratio was around 100 (smallest 0.033, largest 4.1 for the SIV studies and smallest 0.049, largest 3.8 for the SHIV study).

In order to characterize the robustness of the selected treatment effects, we investigated several perturbations to the model and fitting procedure (Tables S6 and S7). We first tested a series models based on biologically-motivated hypotheses (BH) we had formulated about the treatment effects before the model selection was performed. Most of these hypotheses were related to the reservoir reactivation rate, *t_a_*, since the original study interpretations attributed many of the changes in rebound kinetics to effects of immunotherapy on the latent reservoir size. For the SIV data, we tested the study identity (e.g. timing of ART initiation in Study 1 vs 2) influenced *t_a_*, and also whether there was an interaction effect between the TLR7-agonist and the vaccine on *t_a_*. For the SHIV data, we tested whether the antibody or TLR7-agonist influenced *t_a_*, and whether there was an interaction between the TLR7-agonist and the antibody. We next tested a series of alternate models (AM) that were formulated to deal with limitations of practical identifiability that sometimes arose between parameters that could effect rebound kinetics in similar ways. For example, we tested whether the TLR7-agonist treatment effect identified on λ in the SIV data could instead be attributed to *β*, and vice versa for the antibody treatment effect from the SHIV data. We also examined the effect of switching the infered treatment effects on *N_P_* in the SIV data to effects on *m* or *p*, and of switching effects on *m* and *N_P_* in the SHIV data. None of any of these models provided better fits than the one selected by the interative selection process.

As a result of applying our model selection procedure independently to the SIV and SHIV studies, we have inferred that the TLR7-agonist acts on different parameters between the two sets of studies. To ensure that this conclusion was supported by the data, we took the best-fit model inferred in each study and then tested support for that model structure using data from the other study (alternative models beginning in RM in Table S6 and S7). We find that the model structure identified in the SHIV study is strongly disfavored by the SIV data. However, the model structure identified by the combination of SIV studies has reasonable support in the SHIV data relative to the best-fit SHIV model, but the direction of the effects is reversed, making any biological interpretation complicated. Therefore although we still believe the best-fit model is the best interpretation of the treatment effects, we acknowledge that the specific parameters on which the combination off the TLR7-agonist and the PGT121 antibody are acting is less clear in SHIV.

In the statistical model described in Equation S3, we assumed an on-off effect of TLR7 agonist, i.e. *G*(*i, TLR*7) is 0 or 1. We tested several model accounting for the design of TLR7 agonist administration (different number of doses and concentrations administered in subsets of monkeys). The covariate *G*(*i, TLR*7) was replaced by a variable representing the number of doses, the average dose, the cumulative dose or the maximal dose of drug administered (Table S8). None of these alternative models produced a better fit in term of log-likelihood or BIC.

In the SIV data, all of the selected treatment effects for the TLR7-agonist alone occurred only in the context of Study 1 (later ART initiation compared to Study 2) (Table S4). To confirm that this was indeed the best model, we investigated if the effect of the interaction between study identity and TLR7-agonist could be replaced by an TLR7 effect alone. However, such a model exhibited suboptimal log-likelihood and BIC values. We also verified that when restricting the model selection procedure data to only the control groups of animals in each study, the previously identified study effect on *N_P_* remained statistically significant. Finally, we wanted to check that the treatment effects inferred for the TLR7-agonist were not being driven only by the two animals who never rebounded. We repeated the entire model selection procedure for the SIV data but excluding the two non-rebounding animals. No major change in the type and magnitude of the treatment effects was observed.

We tested whether the washout time (*t*_wash_) – a fixed delay between when ART is stopped and when drug has decayed to a low enough level that reactivated virus can seed exponential growth – could be estimated from the data. We found that *t*_wash_ was not practically mutually identifiable with *t_a_*, and profile likelihood suggested that our fixed value of *t*_wash_=3 days was within a range of similarly-likely values that extended upwards by a few days.

#### HIV data

To create a calibrated model of HIV rebound we used data from a collection of individual ACTG studies (5024,5068,5197) in which participants were sampled at least weekly after ART interruption. These studies were previously assembled by Li *et. al*. [55] and included 69 individuals. All subjects in these cohorts originally initiated ART during chronic infection and received no additional immunological interventions along with ART. Median time on ART was ~5 years. At the time of stopping ART, all participants had viral loads less than 50 copies/mL, and ~40% were taking drug combinations involving an NNRTI (EFV or NVP), with the remaining ~60% on non-NNRTI based regimes (NRTI only or PI-based). The original studies were conducted between years 2001 and 2007.

#### HIV model fitting

Overall the strategy used to fit the HIV data was very similar to that used for the SIV/SHIV data. Six model parameters were estimated (λ, *β, t_a_, m, p, N_P_*) while the others were fixed. Fixed parameters values were assumed to be the same between humans and macaques, with the exception of the viral burst size *k*, which was estimated to be 50,000 in macaques [85] but closer to 10,000 in humans [46]; the death rate of actively infected cells *d_I_*, which is estimated to be 1/day in humans [36] vs 0.4/day in macaques [52]; and the concentration equivalent of one actively infected cell in the body *I*_1_, which is 10-fold smaller due to the ~ 10-fold higher body mass of an adult human compared to an adult macaque (~ 7.5 kg vs 75 kg).

The strategy of model fitting as described above for the SIV/SHIV data was applied to the HIV data, except that the only “treatment effect” we attempted to estimate was the impact of NNRTI vs non-NNRTI based ART taken before interruption. To ensure practical identifiability, similarly as before, we fixed the random effect on *p*: otherwise the standard error of parameter p was not correctly estimated. We hypothesized that NNRTI-based therapy may increase the washout time of therapy (before which reactivating latent cells cannot seed growing infection), since NNRTIs are known to have long half-lives compared to PIs and many NRTIs. Previous work observed a difference in median time to first detectable viral load between these groups of participants [55]. In the best-fit selected model, we found that NNRTI-based ART increased the washout time by ~ 3 days (*t*_wash_=3 vs *t*_wash_=6), but did not effect any other viral dynamic parameter. Note that if the time to first detectable viral load is compared between these treatment groups, the rebound delay is much longer, at 8 days (median 16 vs 24 days to rebound), but this is only due to the interval censored nature of the data (due to infrequent sampling). NNRTI-treated individuals had median log_10_ viral loads of 3.0 at first detection vs only 2.4 in non-NNRTI-treated subjects, implying that it took longer to detect their rebound after it crossed the detection threshold. These results highlight the value to fitting entire rebound trajectories instead of examining only the time of first detectable viral load.

#### HIV immunotherapy simulations

We performed simulations to evaluate the possible effects of TLR7-agonist, Ad26/MVA therapeutic vaccine and PGT121 monoclonal antibody treatment in humans. We fit the viral rebound model from Equation (S1) to HIV rebound data, resulting in a posterior distribution of parameters given these data. To generate parameters for a simulated individual, we first sampled a set of model parameters from this posterior distribution. Next, using the population variability and parameter means obtained from this sampling procedure, we sampled the simulated individual’s parameters. Therefore, this sampling procedure captures both the uncertainty regarding paramaters values and population variability. For each treatment scenario, we apply relevant treatment effect on the parameters selected from the model fiting to SIV or SHIV data. The combination of treatment effects used in each hypothetical treatment scenario are described in Tables S10–S14. Using the resulting parameter values, we simulated the trajectory of each simulated individual’s viral rebound after treatment interruption using the package *dslode* in R. It provides us with counterfactual trajectories under treatment of each individual viral rebound assuming that the treatment effect is similar in macaques and humans.

### Derivation: Re-parameterizing the model to account for stochastic reactivation from latency

#### Motivation

We have expressed our model for the dynamics of active and latently infected cells, free virus, and anti-viral immune responses as a system of differential equations (Eqs. (S1)), which implicitly assumes that each variable and parameter is large enough that transitions happen continuously and fluctuations are small relative to the expected dynamics. This is in general a good assumption, especially in the regime we fit to where viral loads are above a detection limit of 50-200 copies/mL. However, there is one reaction for which we believe this assumption often fails: the reactivation of latently infected cells. In the model presented in the main text, we have assumed that reactivation occurs at a continual rate *a*. In reality, reactivations are discrete events occurring to single cells, and when the latent reservoir size is small enough or the per-cell reactivation rate low enough, there could be long waiting times between these events. Previous studies in HIV-infected humans and SIV-infected macaques have estimated these reactivations rates to be between 0.1-5 cells/day [47, 48, 56], and so in the presence of reservoir-reducing therapies, these rates could be much lower. In our data for SHIV-infected macaques especially (Figure 1, Study 3), we often see long delays to rebound followed by relatively rapid viral growth, which are suggestive of low rates of reservoir reactivation.

The differential equation model in Eqs.(S1) can always still be fit to data in which reservoir reactivation happened after a delay, and would just result in a smaller effective *a* value. However, there would be two major problems in interpreting these fit values. One would be that it would not be possible to compare *a* values between two animals or treatment groups and claim that the differences were proportional to differences in reservoir size. As we will show below, when reactivation is common, the inferred *a* value is linearly related to the frequency of reactivation, whereas when reactivation is rare, it is log(*a*) which is proportional to reactivation rate. Another problem is that the variance between individuals in the inferred a value is expected to increase dramatically when reactivation is rare, since the combination of inter-individual variation in reservoir size and the stochastic waiting time until the first reactivation will contribute to the observed time of rebound. One solution could be to fit our data to one of the fully or fully or partially-stochastic models for viral rebound that have been developed previously (e.g. [47, 48, 97]), however, these models are not amenable to the statistically-rigorous group level fitting approaches we wish to employ here. Therefore, as detailed below, we develop a parameter transformation approach that allows us to capture the expected rebound kinetics for any rate of reservoir reactivation. The main idea of this approach is to replace the continuous rate a with a variable *t_a_* which describes the average time between reactivation events for latently infected cells.

Throughout this derivation, we will consider a model of only a single varible - actively infected cells. This is an approximation for the regime where target cells are not yet limited, an effective immune response has not yet kicked in, and free virus is proportional to infected cells. At the end, we will incorporate the results with the full model.

#### Rebound kinetics in the limit of rare reactivation

When latent cells reactivate rarely, the reactivation process can be well described consisting of first a waiting time *t_a_* until the first latent cell reactivates and produces an instantaneous jump in infected cell count to level *I*_1_ (concentration equivalent of 1 infected cell), followed by growth (e.g. [48]). If infected cells release virus and infect new target cells at combined rate *b*, and die at rate *d*, then the differential equation for this process is

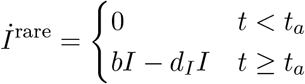

The solution to this equation is

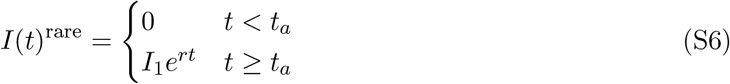

where we define *R*_0_ = *b*/*d_I_* (the basic reproductive ratio) and *r* = *d_I_*(*R*_0_ – 1) (the asymptotic growth rate in the absence of reservoir reactivation, target cell limitation, or immune responses).

The time until rebound, defined as *I*(*t*) = *I_r_*, where *I_r_* is the concentration of infected cells when infection is detectable, can be solved as

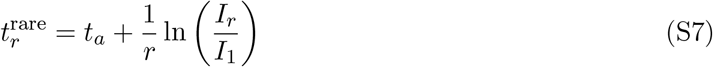

#### Rebound kinetics in the limit of frequent reactivation

When latent cells reactivate frequently, the reactivation process is well described as a continuous rate, *α*, at which cells exit the latent reservoir (e.g. [46, 57]). If each cell contributes a concentration equivalent of *I*_1_, then the dynamics follow a single differential equation

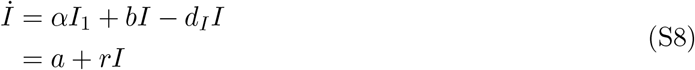

were we define *α* = *αI*_1_ (the concentration of latent cells exiting the reservoir per day), and as before, *R*_0_ = *b*/*d_I_* and *r* = *d_I_*(*R*_0_ – 1). This equation has the solution, for all *t* > 0

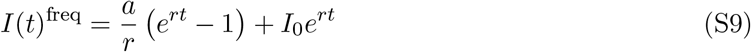

The time until rebound, defined as *I*(*t*) = *I_r_*, can be solved as

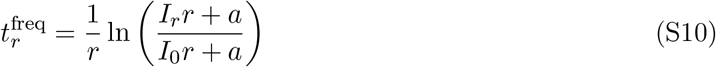

The initial condition, *I*_0_, is the equilibrium value of Eq. (S8) with *b* = 0, which gives the value of *I* during administration of ART and therefore at the time of ART stop

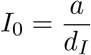

which results in the solution

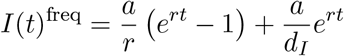

and the updated rebound time

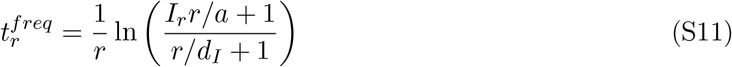

#### Discontinuity between the two regimes

First, we will show that there is a discontinuity between these regimes in terms of the time to rebound as a function of the amount of reactivation. To compare them, first note that if cells exit the latent reservoir at rate *a*, and these events are independent and time homogeneous, then the average time between reactivation events is

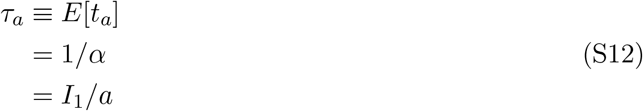

and the distribution of individual waiting times *t_a_* follows *p*(*t_a_*) = (1/*τ_α_*)*e*^−*t_a_*/*τ_a_*^.

If we look at the limit of rare reactivation dynamics, then the average time to rebound, for a given average waiting time *τ_a_* between reactivation events, is

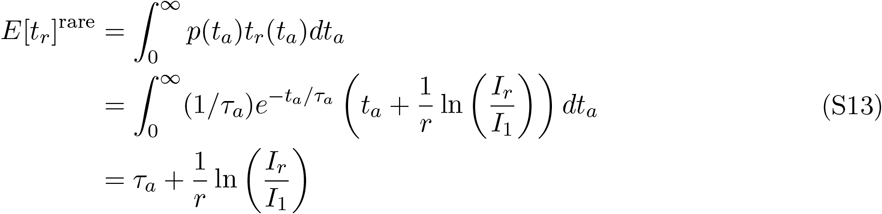

which can similarly be expressed in terms of the latent reservoir reactivation rate *a* using Eq. (S12)

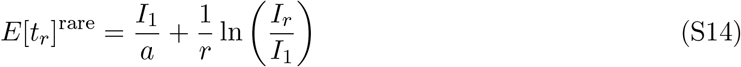

Next, we look at the formula for frequent reactivation dynamics, and see what happens if reactivation becomes rarer. Does it approach the expression for the rare reactivation regime, Eq. (S14)? Taking the limit of Eq. (S11) as a approaches zero, we get

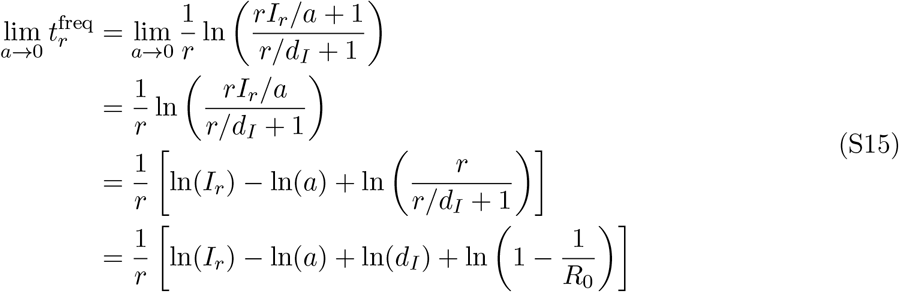

which can similarly be expressed in terms of the average time between latent cell reactivates, by setting *a* = *I*_1_/*τ_a_* using Eq. (S12)

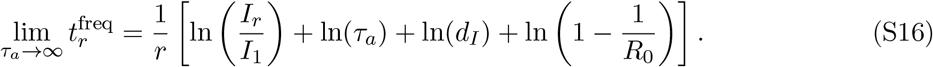

We can see that Equation (S13) and Equation (S16) do not match. Rebound time should grow linearly with *τ_a_* in the rare reactivation regime (Eq. (S13)) but in Eq. (S16) it only grows logarithmically. Generally, Eq. (S16) underestimates the rebound time, since *I_r_* ≫ *I*_1_ and *R*_0_ > 1.

We can understand qualitatively why the two models don’t match. The rare reactivation model assumes that even in the case of instantaneous reactivation (*a* → ∞ or *τ_a_* → 0), the infection only starts growing from a level *I*_1_. Only a single reactivating cell contributes to rebound. However, in the frequent reactivation model, if reactivation is high then the initial condition is much larger than *I*_1_, since there would have been many reactivated cells around prior to ART stop which can begin growing immediately upon ART stop. And, cells can continue reactivating during rebound, which further increases the rate at which infection grows beyond just at rate *r*.

**Figure S1:**
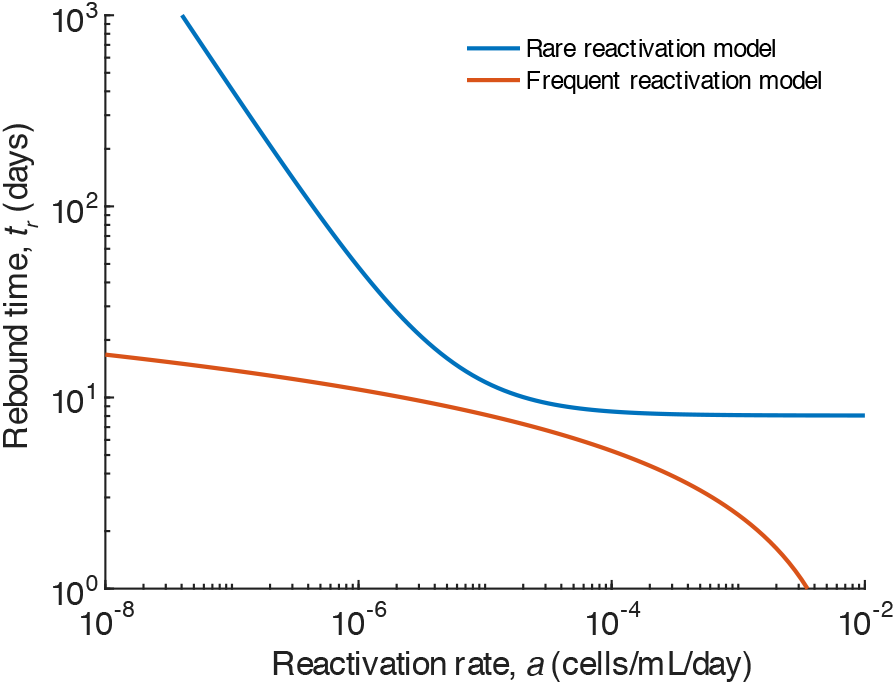
Comparison of expected rebound times under rare and frequent reactivation models. Expected time to rebound (to a viral load of 50 copies/mL) according to the simplified models in Equations (S14) (rare, red) and (S11) (frequent, blue) for a wide range of latent cell reactivation rates. When reactivation is very rare (small *a*), the frequent reactivation rate model (blue) dramatically underestimates the expected time until rebound. For very frequent reactivation (large *a*), the rare reactivation rate model (red) fails to predict almost instantaneous rebound, primarily because it fails to account for the correct initial conditions. Both models use the following parameters which are estimated for macaques: *I*_1_ = 4 × 10^−5^ cells/mL, *I_r_* = 50 copies/mL/(*k/c*)=0.025 cells/mL (with *k* = 50,000 virions/cell/day and *c* = 25), *r* = 0.8 day^−1^, *d_I_* = 0.4 day^−1^. Sources for these parameters are given in Tables 1 and S2.

#### Bridging the two regimes

We can bridge the two regimes by thinking of a model that would apply in the regime of intermediate *α*, where reactivation occurs relatively quickly and a few reactivation events contribute to rebound. Assume that cells reactivate exactly every Δ time steps. This derivation is inspired by a calculation by Pinkevych *et al* [98]. Each cell that reactivates starts at level *I*_1_ and then grows, according to Eq. (S6), exponentially at rate *r*. This gives a formula for the size of the total infection

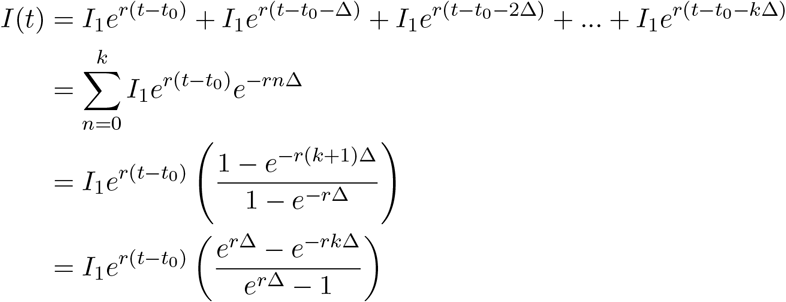

where *t*_0_ is the time of the first reactivation, and *k* + 1 is the number of reactivations that happen before time *t*. It is the highest integer such that *t* – *t*_0_ – *k*Δ ≥ 0, which implies that (*t* – *t*_0_)/Δ – 1 < *k* ≤ (*t* – *t*_0_)/Δ. We set *t*_0_ = Δ = *t_a_*, because we want *t_a_* to have the interpretation of being the time of the first reactivation and will assume that it is also representative of the average waiting time. We choose the highest possible *k*, so *k* = (*t* – *t*_0_)/Δ = *t*/*t_a_* – 1.

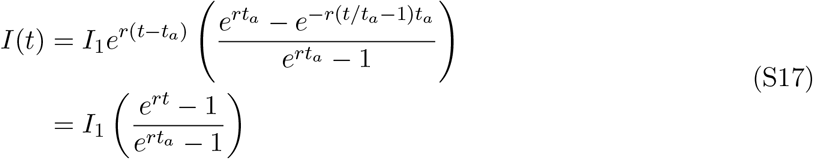

Compare this equation to our previous expression for continuous reactivations, Eq. (S9). They are equivalent when

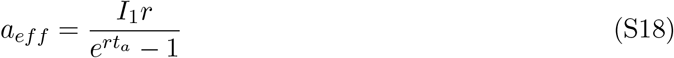

and

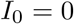

Thus, conceptually, this method for including multiple reactivations has made our formula for rare reactivations (Eq. (S6)) closer to the one for frequent reactivations (Eq. (S9)) by accounting for the contributions of multiple reactivating cells. However, it would still underestimate rebound time when reactivation is really common because it still assumes the initial cell level (*I*_0_) at the time of ART interruption is zero.

Instead, we want to choose a value of the initial condition *I*_0_ which is a function of *t_a_* and makes the new model in Eq. (S17) match in all regimes. When *t_a_* is large, we want *I*_0_ = 0. When *t_a_* is small, we want *I*_0_ = *a*/*d_I_* = *I*_1_/(*d_I_t_a_*). One option is

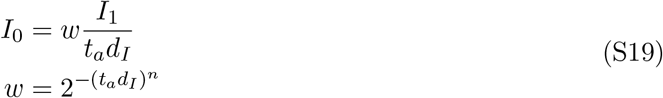

This function describes a sigmoidal curve that goes from one to zero, switching at *t_a_* = 1/*d_I_*. The constant *n* controls the sharpness of the interpolation (higher *n*, sharper transition).

The rational behind this function is that when *t_a_d_I_* ≪ 1, there are many cells reactivated even at the moment ART is stopped (at average level *I*_1_/(*d_I_t_a_*)), while when *t_a_d_I_* ≫ 1, there are usually no latent cells activated at the time ART is stopped. In reality, the number of cells present at ART stop is a random variable (Poisson-distributed with mean *I*_1_/(*d_I_t_a_*), see [47]), which a deterministic model cannot incorporate exactly. However, this is a reasonable approximation to the average behavior and reproduces the average rebound time when *t_a_* is the mean time between reactivations.

Note: With this new *I*_0_, it will not be true that *I*(*t_a_*) = *I*_0_. As soon as there is a non-zero initial condition, there will be some growth that happens before the first post-ART reactivation. This growth is due to cells that reactivated before ART stopped. Therefore, *I*(*t_a_*) is equal to *I*_0_ plus this older growth.

We can analytically calculate the rebound time for this model. Using Eq. (S10) along with Eq. (S18) and Eq. (S19), gives

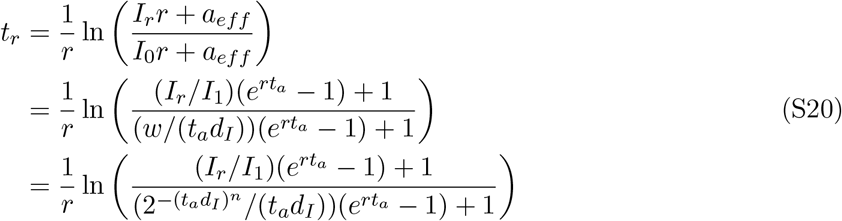

This equation could similarly be expressed as a function of the latent reservoir reactivation rate *a* using Eq. (S12)

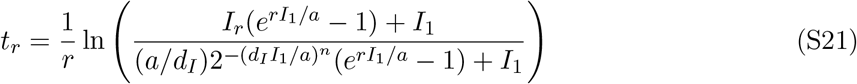

We can check that in the limit of rare reactivation (large *t_a_*), the rebound time will approach the value for the original rare reactivation model (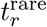, Eq (S7)):

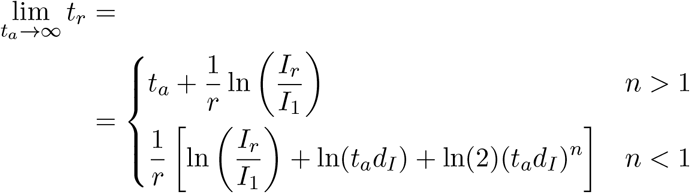

The first expression, with *n* > 1, gives the correct limit. However, if *n* is too small (less than one), then the weighing function *w* doesn’t decay fast enough with *t_a_*, and so the initial condition *I*_0_ doesn’t decay to zero fast enough, and the time to rebound is underestimated (rebound happens too fast). Using *n* = 3 gives a smooth function for rebound times versus reactivation rate with biological values of other parameters and gives the correct behavior in the limit of rare reactivation (large *t_a_*) (Figure S2).

**Figure S2:**
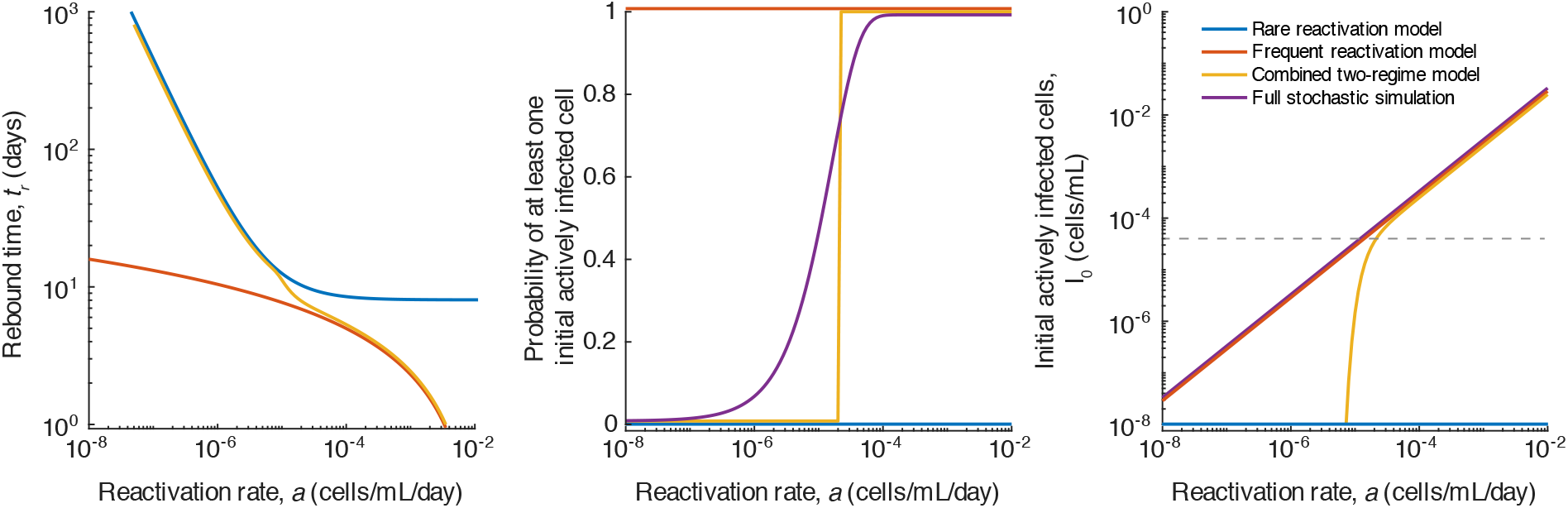
The combined approach matches expected rebound times under rare and frequent reactivation models. A) Expected time to rebound (to a viral load of 50 copies/mL) according to the simplified models in Equations (S14) (rare, red) and (S11) (frequent, blue), and the combined model in Eq. (S21), for a wide range of latent cell reactivation rates. The combined approach agrees with the rare reactivation model for low reactivation rates and the frequent reactivation model for high reactivation rates. B) The probability that at there is at least one actively infected cell present at the time of ART stop. This is zero in the rare reactivation model, one in the frequent reactivation model, and interpolates between them in the combined model. In a fully-stochastic model, the initial value is expected to be a Poisson-distributed random number with mean *a*/*d_I_*. C) The number of actively infected cells present at the time of ART stop. In the rare reactivation model, this is always zero, whereas in the frequent reactivation model, this quantity increases linearly with the reactivation rate, and equals the expected value of the fully-stochastic model. In the combined model, this value approaches zero more rapidly to account for the fact that in reality the expected value is often below the level corresponding to a single cell (dashed line). All models use the following parameters which are estimated for macaques: *I*_1_ = 4 × 10^−5^ cells/mL, *I_r_* = 50 copies/mL/(*k/c*)=0.025 cells/mL (with *k* = 50, 000 virions/cell/day and *c* = 25), *r* = 0.8 day^−1^, *d_I_* = 0.4 day^−1^, *n* = 3. Sources for these parameters are given in Tables 1 and S2.

We can put all of this together to define a model that works in both regimes:

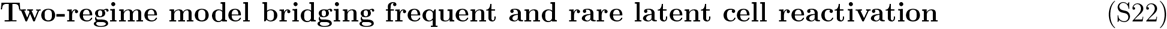

***Functions to connect stochastic and deterministic regimes:***

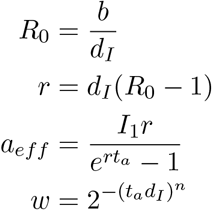

***Initial conditions:***

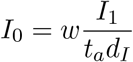

***Equations:***

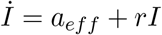

Note that these equations work even when *R*_0_ < 1 (*r* < 0), and *a_eff_* = *I*_1_ when *R*_0_ = 1 (*r* = 0) (although numerically it may be undefined).

#### Conditioning on survival reactivating cells in the stochastic regime

We previously analyzed the dynamics of rebound assuming that cells reactivate from latency every *t_a_* days (i.e. at rate 1/*t_a_*) and that infection then grows exponentially towards rebound. This model is a simplification, since in reality infection dynamics are a fully stochastic process. Developing a model to track every stochastic reaction between a cell and virus is beyond the scope of this work, and such a model would not be identifiable from typical *in vivo* measurements of viral kinetics. However, recognizing the underlying stochastic nature of these dynamics leads to an important correction to our work.

While some reactivating latent cells will produce a chain of infection that eventually leads to rebound, others will - simply by chance - end up going extinct. Without specifying any details of the underlying stochastic processes, we can define the probability of long-term survival of the infection started from a single reactivating cell (often called the “establishment” or “survival” probability) as *p*_surv_ ∈ [0,1]. Then the average time between *surviving* reactivations is *t_a_*/*p*_surv_.

While overall, the expected dynamics averaged over all reactivating cells is described by the deterministic equations,

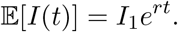

If we condition on survival of the reactivated cell, then the expected dynamics are larger by a factor of 1/*p*_surv_:

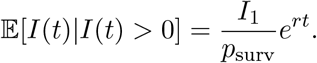

Together, this means that Eqs. (S22) can be updated with an effectively longer interval between reactivating latent cells (*t_a_* → *t_a_*/*p*_surv_) and an effectively higher initial concentration of actively infected cells (*I*_1_ → *I*_1_/*p*_surv_).

The value of the survival probability and its relationship to the other model parameters depends on the details of the underlying stochastic process. In all cases, *p*_surv_ will be higher when *R*_0_ is higher. As an example, we consider the relatively generic stochastic process used to describe reactivation from latent infection in Hill et al [47]:

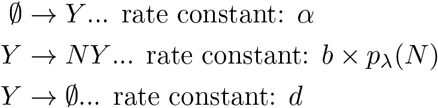

This model tracks only actively infected cells. In this notation, *Y* represents an individual cells and Ø represents no cells, and the arrows represent events that change the number of cells. Mathematically this process is a type of burst-death-immigration branching process. A reactivation event from latency produces an actively infected cell at rate *α*, where *t_a_* = 1/*α*. This cell can either die (at rate *d*) or produce a collection of virions (at rate *b*) that results in the infection of *N* other cells, where *N* is a Poisson-distributed random variable with parameter λ, *p*_Λ_(*N*) = (*e*^−Λ^λ^*N*^)/(*N*!). After an infection event, the original cell dies.

The overall survival probability for a single reactivated cell is the weighted sum of the probability of producing *N* offspring and the probability that at least one of these offspring survives.

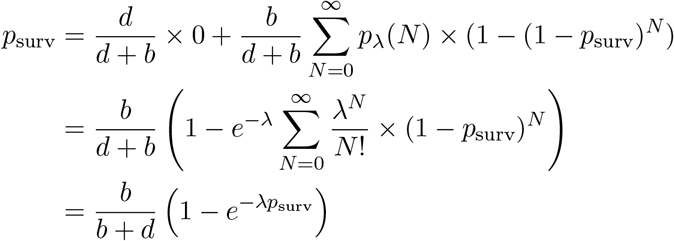

Since it is generally impossible to measure the underlying stochastic parameters (e.g. *b, d*, and λ), we want to replace them in this equation with composite parameters that are more accessible. We use the basic reproductive ratio, 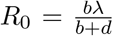, which describes the average number of secondary infections arising from a single cell, and *ρ*, the so-called Fano factor, which is the ratio of the variance to the average for the same quantity (*ρ* = 1 + *d*λ/(*b* + *d*)). Previous studies have suggested that *ρ* ~ 10 (reviewed in [47]). Then, we find that the survival probability satisfies the implicit condition

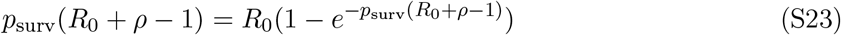

Note that the survival probability is only non-zero when *R*_0_ > 1. Otherwise extinction is guaranteed.

The solution to this implicit definition can be expressed in terms of the Lambert W function:

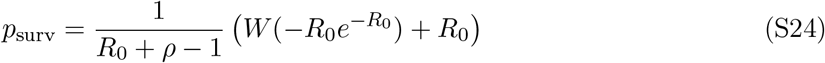

Conditioning on survival for reactivating lineages in the stochastic regime leads to the following combined model. Note that for the *R*_0_ < 1 case *p*_surv_ = 0 and we have taken the limit of expressions as *p*_surv_ → 0 to avoid undefined expressions.

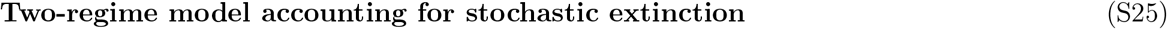

***Functions to connect stochastic and deterministic regimes:***

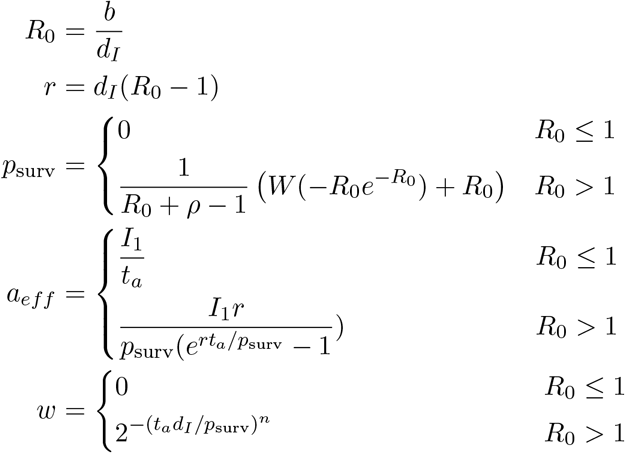

***Initial conditions:***

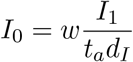

***Equations:***

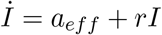

#### Accounting for numerical errors

For all differential equation solvers we have tested (e.g. R, Matlab, Monolix), numerical errors often arise when evaluating Eqs. (S22) or (S25) when *t_a_* is large. This occurs due to the extremely small values of *a_eff_* and *w* (and therefore *I*_0_) and the extremely small values of *I*(*t*) inferred for *t* < *t_a_* with this continuous approximation. We can get around this by choosing not to start integration for *I*(*t*) right at *t* = 0 if both *a_eff_* and *I*_0_ are very small, but instead, choosing a time *t_go_* such that at least one of them is larger. We also want to make sure this time is not too large, because then, infection levels could have grown out of the simple regime we are considering now (no target cell limitation, no immune control).

An easy solution is to choose a *t_go_* such that *I*(*t_go_*) = *I*_1_. Since *I*_1_ is the true biological minimum of infection, we know that for *I*(*t*) < *I*_1_ any approximations about exponential viral growth are valid.

**Figure S3:**
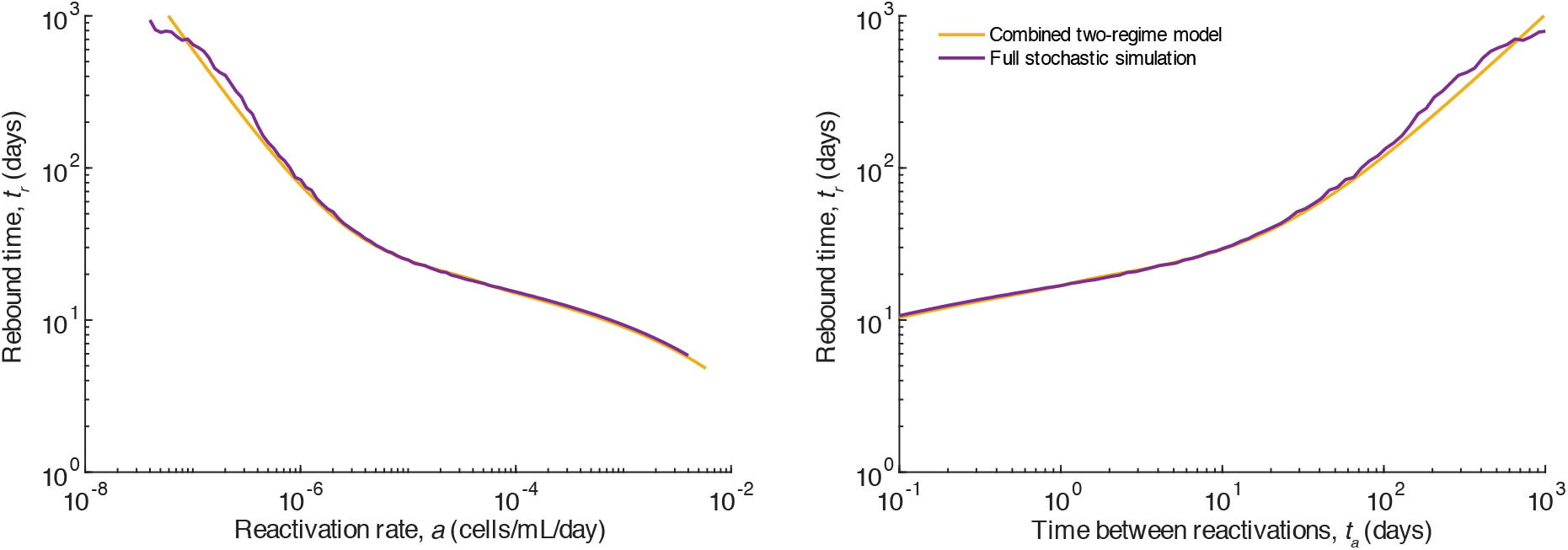
The combined approach matches the results of a fully-stochastic simulation. A) Expected time to rebound (to a viral load of 50 copies/mL) according to the combined model in Eqs. (S22) for a wide range of latent cell reactivation rates. The fully-stochastic simulation was conducted using the code from [47] but without any death of latently infected cells (since reservoir decay is not included in our models here, and is likely not relevant on the timescales in these studies). B) Same but plotted versus the time between latent cell reactivations. All models use the following parameters which are estimated for humans: *I*_1_ = 4 × 10^−6^ cells/mL, *I_r_* = 50 copies/mL/(*k/c*)=0.125 cells/mL (with *k* = 10,000 virions/cell/day and *c* = 25), *r* = 0.4 day^−1^, *d_I_* = 1 day^−1^, *n* = 3. Sources for these parameters are given in the main text methods describing the human simulations and S2.

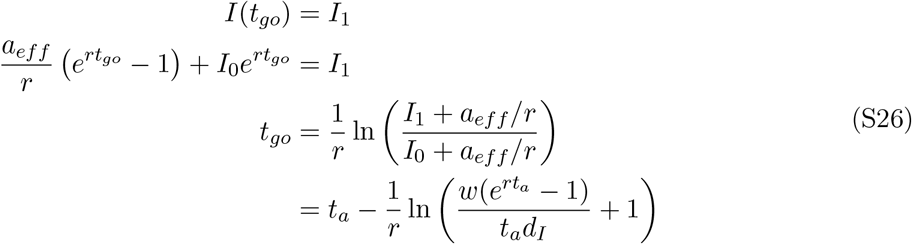

Then we integrate the equation

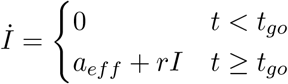

with initial condition

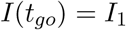

which has the solution

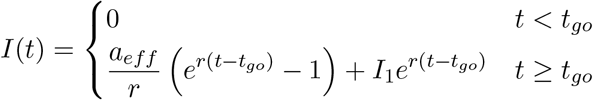

Note that sometimes the reservoir reactivation rate will be high enough that more than one reactivated cell is expected to be present at the time of ART interruption, on average, i.e, *I*(0) = *wI*_1_/(*t_a_d_I_*) > *I*_1_ (from Eq. S25). The formula in Eq. (S26) would give a negative *t_go_* in this regime. In this case, the equation can be integrated immediately from time zero onwards without numerical errors. Hence we set *t_go_* = 0 and use the initial condition of Eq. (S25).

When we combine these corrections for numerical errors with the corrections to account for stochastic extinction (Eqs. (S25)), e.g. take *t_a_* → *t_a_*/*p*_surv_ and *I*_1_ → *I*_1_/*p*_surv_, this leads to the updated model:

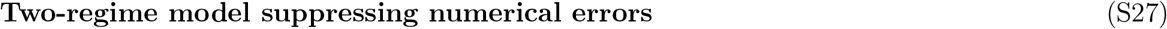

***Functions to connect stochastic and deterministic regimes:***

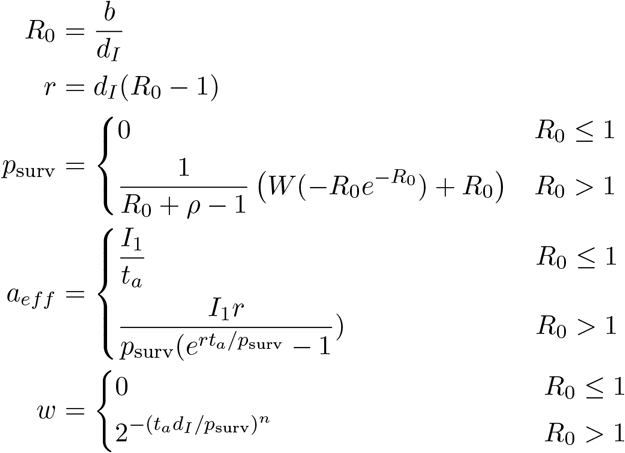

***Functions to avoid small number errors:***

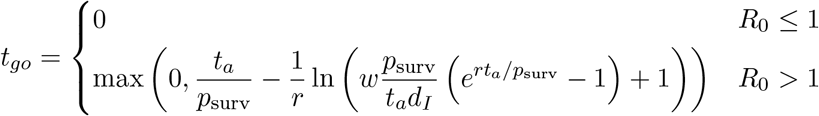

***Initial conditions:***

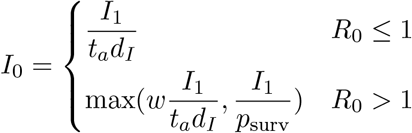

***Equations:*** For all *t* > *t_go_*

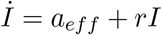

#### Calculating the long-term growth rate when free virus is included

The model we have shown tracks only infected cells, as well as ignoring the immune response and target cell limitation. The final model can be augmented to includes these effects by simply adding the extra variables and terms to the system of equation for *I*(*t*). However, in doing so, we must make a slight alteration in our expression for *r*, the early growth rate of infection (before target cell limitation or immune response has set in). Previously, we had used the expression *r* = *d_I_*(*R*_0_ – 1), but this is not valid when we track free virus as well as infected cells.

Consider a set of viral dynamics equations tracking infected cells along with free virus:

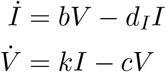

Which can be expressed in matrix form, with 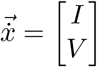, as

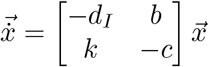

The eigenvalues λ of this matrix satisfy the polynomial

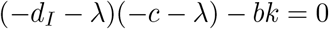

Which has the solutions

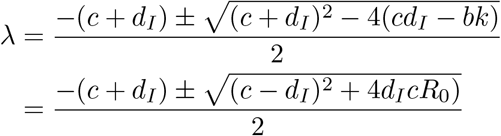

The positive root of this equation dominates the solution in the long term, so we set the parameter *r* in our model to be

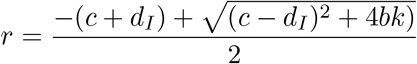

In this model, the basic reproductive ratio is *R*_0_ = *bk*/(*cd_I_*), and so *r* can be expressed in terms of *R*_0_ as

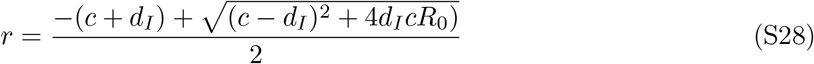

If *R*_0_ > 1, then *r* > 0. This equation can be re-arranged for *R*_0_, as

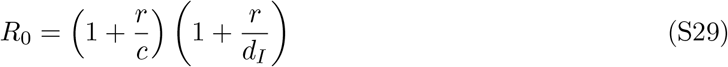

which reduces to the simplified formula, *R*_0_ = *r*/*d_I_* + 1 or *r* = *d_I_*(*R*_0_ – 1) only if *r* ≪ *c*. While estimates of *R*_0_ and *r* using the simplified formulas will only be off by around 5% for values of *c* = 23 /day and *r* ~ 1/day observed in this study, this will significantly alter our estimates of *a_eff_* (Eq. S22) and lead to biased estimates of other parameter values.

#### Incorporating the antiviral immune response

The methods we have described above for modeling stochastic reactivation from latency in the context of the deterministic model of viral dynamics expressed in Eq. (S1) involve considering events that happen before or near the time of the first reactivated latent cell. Before this, we assume that if there is any immune response present, it is at the steady state level that would be stimulated if viral load and infected cells were at the average values expected during ART. The presence of immunity pre-ART acts to decrease *R*_0_ and *r*.

During ART, the expected average size of the infected cell and viral population are

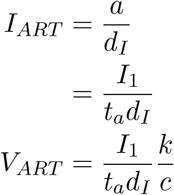

Target cells are at their pre-infection equilibrium level

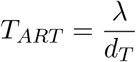

And the steady state level of immune precursors and effectors are

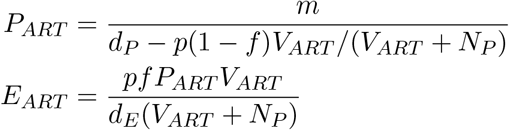

The expression for the basic reproductive ratio, *R*_0_, is thus

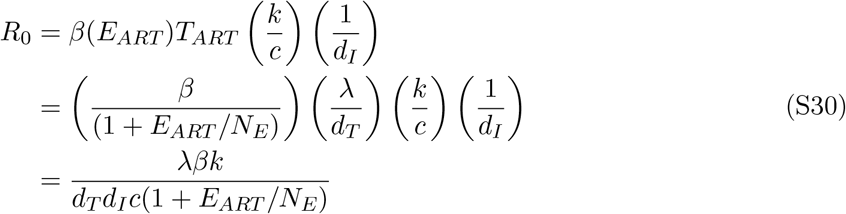

Note that it is theoretically possible in this model to be in a parameter regime where the immune response grows indefinitely during ART even when the infection is of a constant size, if that size is large enough (e.g. *Part* and *Eart* undefined). However, this never occurs for realistic values of the model parameters, but during a parameter fitting procedure in which a wide range of (non-realistic) values may be sampled at some point, it is wise to create a conditional expression to avoid such situations. We do so by setting *Part* = 1000*N_E_* (Since *N_E_* is mutually non-identifiable with *m*, the value chosen for it determines the scale of *m* and therefore also *P* and *E*).

With these expressiosn for *R*_0_ and *r*, the final model that we fit to the data is:

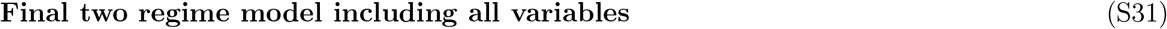

***Variables:*** (Observed) *V*, (Unobserved) *T, I, P, E*

***Basic parameters:*** (Fit) λ, *β, t_a_, N_P_, p, m*, (Fixed) *d_T_, d_I_, d_P_, d_E_, c, k, N_E_, f*, time for antiretroviral therapy to wash out (*t*_wash_), concentration equivalent of one infected cell in body (*I*_1_), mean-to-variance ratio for virus production (*ρ*), and interpolation constant (*n*).

***Time-averaged values of variables during ART (**β = 0)*:

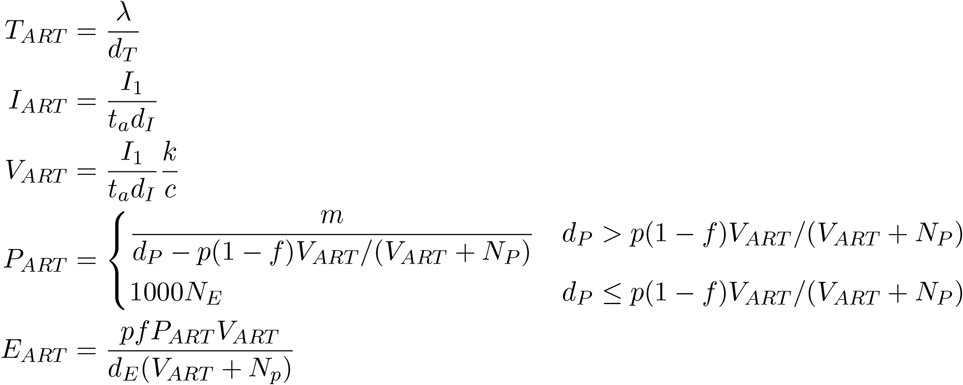

***Functions to connect stochastic and deterministic regimes:***

Basic reproductive ratio:

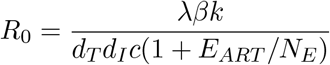

Early exponential growth rate:

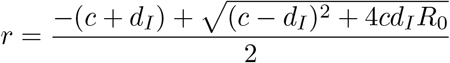

Survival probability starting from single actively-infected cell:

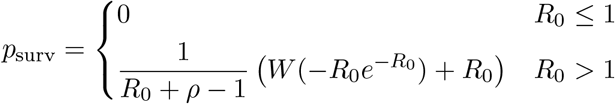

Effective reactivation rate of latently-infected cells:

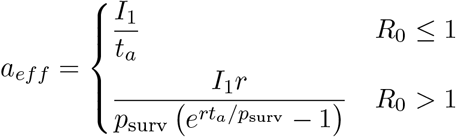

Interpolation function:

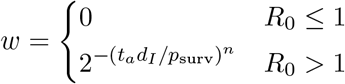

***Functions to avoid small number errors:***

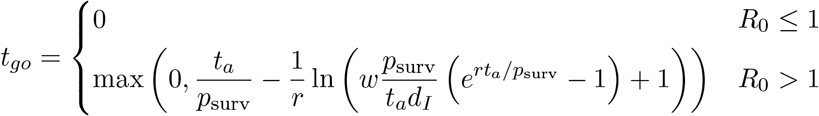

***Initial conditions:***

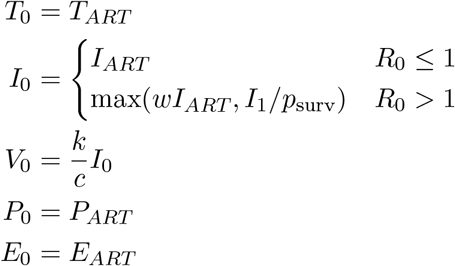

***Equations:*** For all *t* > *t*_wash_ + *t_go_*

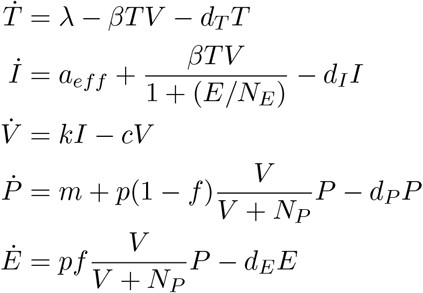

#### Parameters involved in stochastic-to-deterministic model

In order to use our simple ordinary differential equation model (Eq. (S1)) to simulate SIV/HIV dynamics in both the deterministic and stochastic regimes using the *t_a_* → *a* transformations described in Eqs. (S31), we needed to introduce four additional fixed parameters that occur in the transformation. Here we desrcibe the source of the value we used for each.

##### Concentration equivalent of one infected cell (*I*_1_)

Our differential equations (Eq. (S1)) track the levels of virus and infected cells as concentrations per mL of blood (Table S1). However, in the regime where reactivation from latency is rare, what matters is the time until the first individual cell reactivates. Therefore, we need to be able to scale between infected cell concentration and total body numbers.

We assume that both virus and CD4+ T cells are well mixed throughout the blood and lymph tissue, but that the concentrations are higher in the lymph tissue by a fixed fraction. For an adult human weighing 75 kg, there are around 3L of blood plasma, since total body water is ≈ 60% of weight, of which 1/3 is extracellular and 1/5 of that is plasma (e.g. overall 4% of weight is plasma) [99]. Plasma makes up abut 60% of blood, for a total of 5L of blood [99]. Estimates suggest that only around 2% of lymphocytes are circulating at any time [100], so there are 50 cells in the lymph tissue for every one in the blood. Consequently, for humans, the blood concentration equivalent of one infected cell anywhere in the body is *I*_1_ = 1 cells/(5,000 mL×50)=4 × 10^−6^. We assume this is the level that infection starts from after the first fated-to-survive cell reactivates from latency.

For macaques, we assume a weight of 7.5 kg but all other ratios the same, corresponding to 0.5 L of blood and *I*_1_ = 4 × 10^−5^.

##### Slope of interpolation function for infected cells at time of ART stop (*n*)

We wanted to use an integer value of *n* > 1 that resulted in smooth interpolation between the regimes of rare vs frequent reactivation of latent cells for approximately realistic parameter values. We found that *n* = 3 worked well (see Figure S2).

##### Variance-to-mean ratio for secondary infections from a single cell (*ρ*)

In any stochastic branching process, such as the spread of virus between infected cells early in infection or rebound, extinction is possible even if on average the process is expected to grow. Thus, the probability that an infection starting from a single reactivated latent cell will eventually lead to viral rebound (the survival probability, *p*_surv_) is generally less than one. In previous work [47], we found that for a relatively general stochastic model of viral dynamics, the survival probability was a function of only the mean number of secondary infections (the basic reproductive ratio, *R*_0_) and the ratio of the variance to the mean (the Fano factor, *ρ*). While *R*_0_ can be estimated from the rate of exponential growth during acute infection or rebound, *ρ* concerns stochastic events that cannot be measured with bulk viral load measurements. Instead, we roughly estimated *ρ* ≈ 10 based a) in vivo measurements of the effective population size of HIV vs the census population size, which can be directly related to variance in secondary infections, b) in vivo easurements of the rate of exit of drug resistant HIV from the latent reservoir, and c) in vitro measurements of the variance in viral gene expression using flourescent reporter proteins. More recently, Hataye *et al*. [101] have conducted a detailed analysis of HIV infection ex vivo at the single cell level and estimated *ρ* ≈ 30, which gives survival probabilities in the same general range.

##### ART washout time (*t*_wash_)

When antiretroviral therapy is stopped, drugs are not instantaneously cleared from the body. Consequently, even if reactivated latent cells are present at the time of ART cessation, rebound cannot occur immediately, since drugs will still be at levels that suppress virus replication for some time. We call the time until drug levels decay such that exponential viral growth can occur (*R*_0_ > 1) the washout time, *t*_wash_. Ignoring washout time in model fitting will lead to underestimates for the rate of reactivation of cells from the latent reservoir (*a, t_a_*), and underestimates for treatment effects that reduce the reservoir.

Theoretically, the rate of decay of drug suppression should be able to be calculated based on the drug half-life, the drug concentration at the time of ART stop, and the dose-response curve for the viral inhibitory effect of the drug [38, 102]. However, these methods can quickly become complex and unreliable due to in vitro-in vivo differences, adjustments for drug protein binding and intracellular pro-drug metabolism, pharmacodynamic interactions between drugs in combination therapy. Instead, we estimated washout time from clinical data. One study of dolutegravir (DTG) washout in HIV+ individuals found that drug levels became non-suppressive between day 3 and 4 in most subjects [103]. Separately, studies of “short-cycle” therapy involving tenofovir and emtracitabine (TFV/FTC) have found that taking 5 days of ART followed by 2 days off [104], or 4 days on + 3 days off [105], is as effective as daily therapy, whereas 7 day on/7 day off [106] or less than 3 doses a week [107] led to more treatment failures. In one study, most patients were on TFV/FTC plus an NNRTI or PI, and drug level monitoring during treatment breaks found subtherapeutic levels of the NNRTIs and PIs, implying that TFV/FTC must be responsible for continued viral suppresson during off days [105] Collectively, this suggests a DTG/TFV/FTC washout time of around 3 days. We assumed *t*_wash_ was the same in macaques and humans, and between SIV and SHIV.

Note that ideally *t*_wash_ should not be an external parameter but be fit as part of a model that incorporates drug decay, since the time until drug is no longer suppressive also depends on the intrinsic fitness of the virus (*R*_0_), which is fit from rebound kinetics. Virus with higher *R*_0_ will be able to begin expanding at higher residual drug levels, making *t*_wash_ smaller. However, in the interest of not further complicating our model, we just chose a fixed value. As described in the **Model Selection** section, *t*_wash_ was not identifiable from the data.

**Table S2:** Parameters used to transform the model between the deterministic and stochastic regimes

|  | Description | Units | Value | Source |
| --- | --- | --- | --- | --- |
| $I_1$ | Concentration equivalent of one infected cell | cells mL<sup>-1</sup> | $4 \times 10^{-5}$ (macaques)<br>$4 \times 10^{-6}$ (humans) | [99, 100] |
| $n$ | Slope of interpolation function for infected cells at time of ART stop | 1 | 3 | |
| $\rho$ | Variance-to-mean ratio for secondary infections from a single cell | 1 | 10 | [47] |
| $t_{\text{wash}}$ | ART washout time | days | 3 | [103–107] |

**Figure S4:**
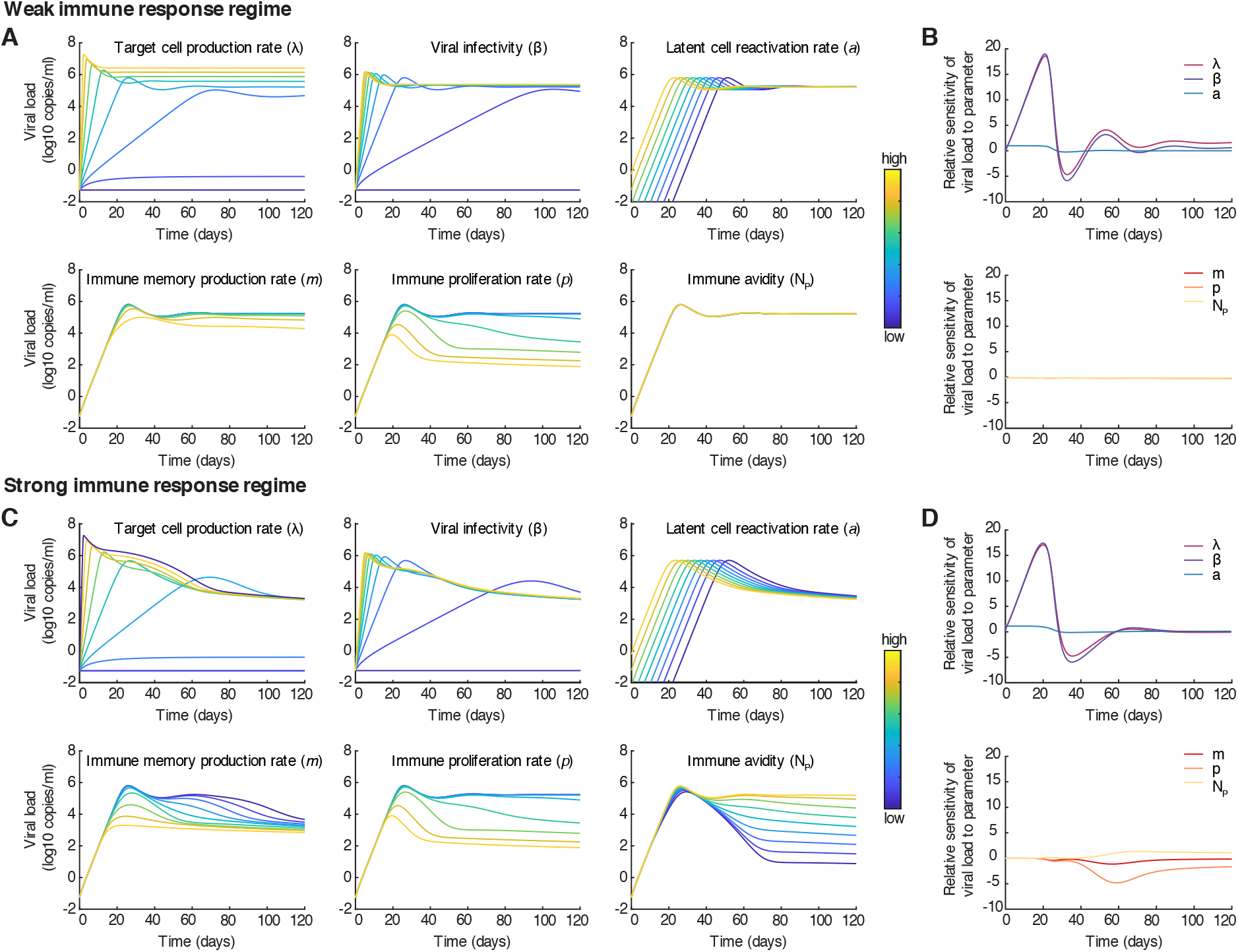
Impact of kinetic parameters on viral rebound trajectories in two regimes. Top: Weak immune response (*p* = 0.1). A) Viral load trajectories produced by the model for different values of the six fitted parameters. B) Sensitivity of viral load to parameter values over time. Relative sensitivity to parameter *θ* is defined as 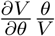. Bottom row: Strong immune response (*p* = 1). C) Same as A. D) Same as B. Parameter values, when not varied and [min,max] when varied: λ = 50 [0, 500] cells mL^−1^ day^−1^, *β* = 5 × 10^−7^ [0, 20] mL copies^−1^ day^−1^, = 10^4^ cells mL^−1^, d_T_ = 0.05 day^−1^, *a* = 10^−5^ [10^−12^, 10^−4^] cells mL^−1^ day^−1^, *d_I_* = 0.4 day^−1^, *k* = 5 × 10^4^ virions cells^−1^ day^−1^, *c* = 23 day^−1^, *m* = 1 [0.01, 100] cells mL^−1^ day^−1^, *f* = 0.9, *p* = 0.1 (A,B) or *p* = 1 [0.01, 10] (C,D) day^−1^, *N_P_* = 10^4^ [10^2^, 10^6^] copies mL^−1^, *d_E_* = 1 day^−1^, *d_P_* = 0.001 day^−1^.

**Figure S5:**
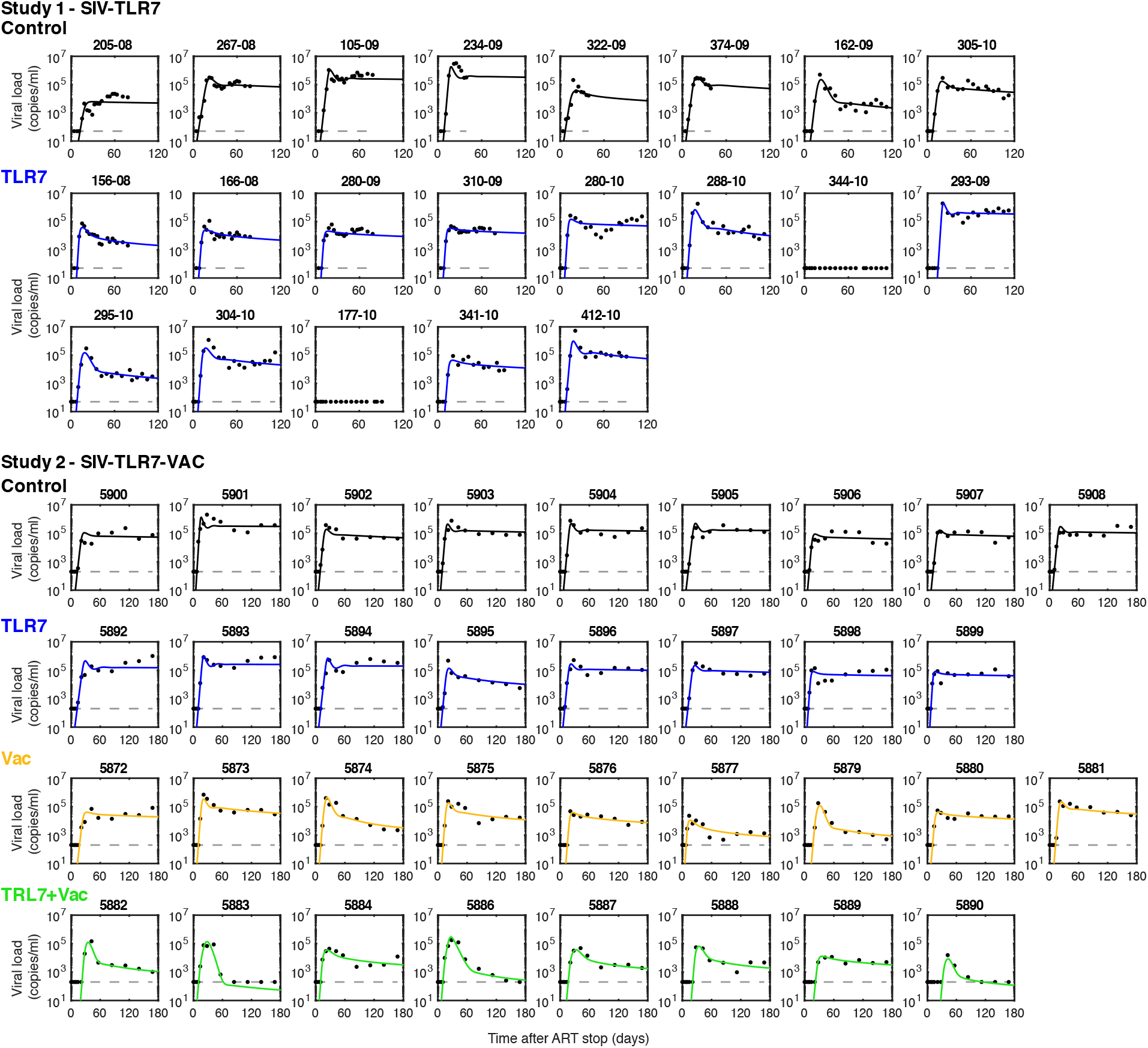
Viral load values and model fits during rebound for all SIV-infected animals in Study 1 and 2.

**Figure S6:**
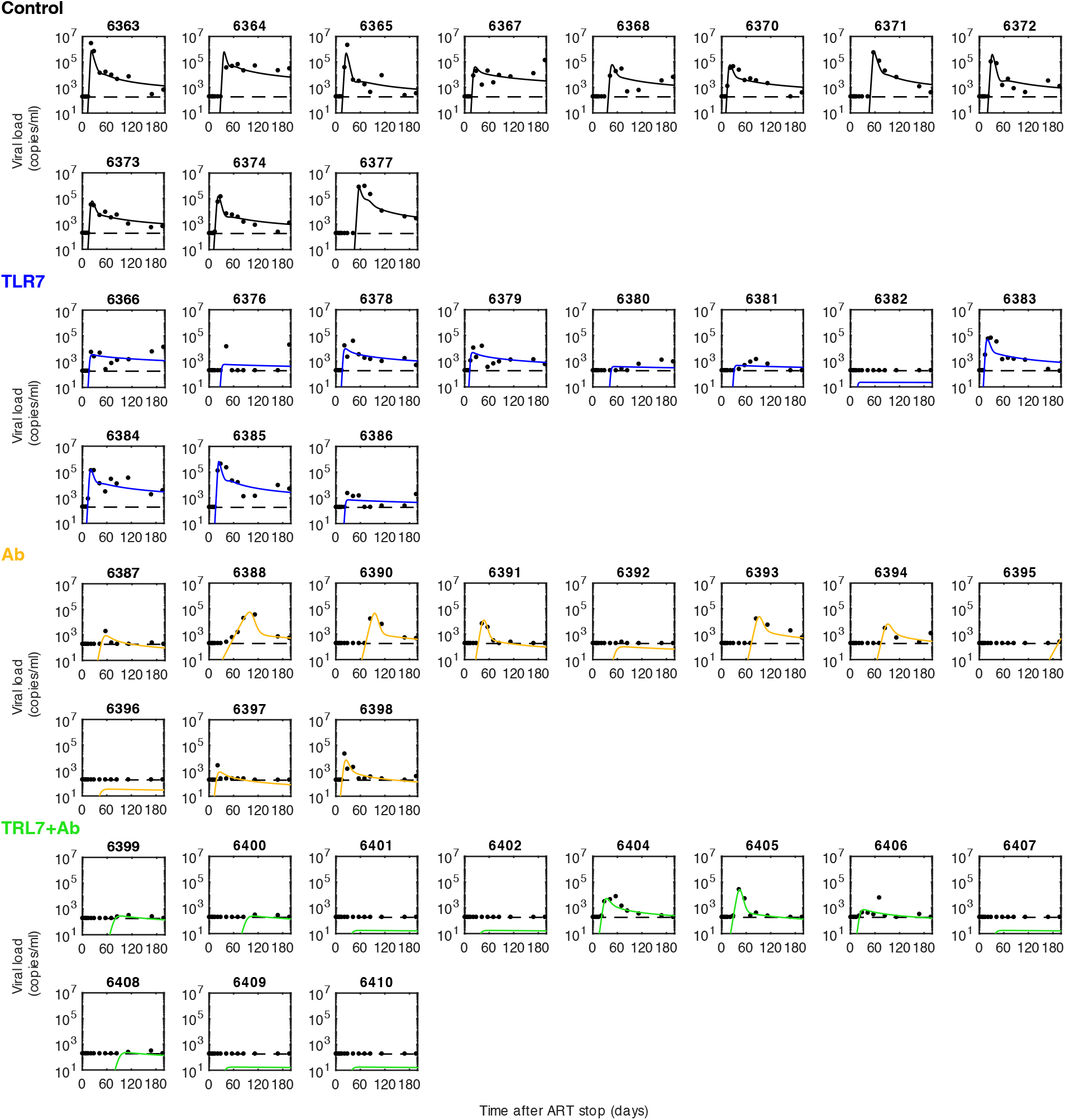
Viral load values and model fits during rebound for all SHIV-infected animals in Study 3.

**Figure S7:**
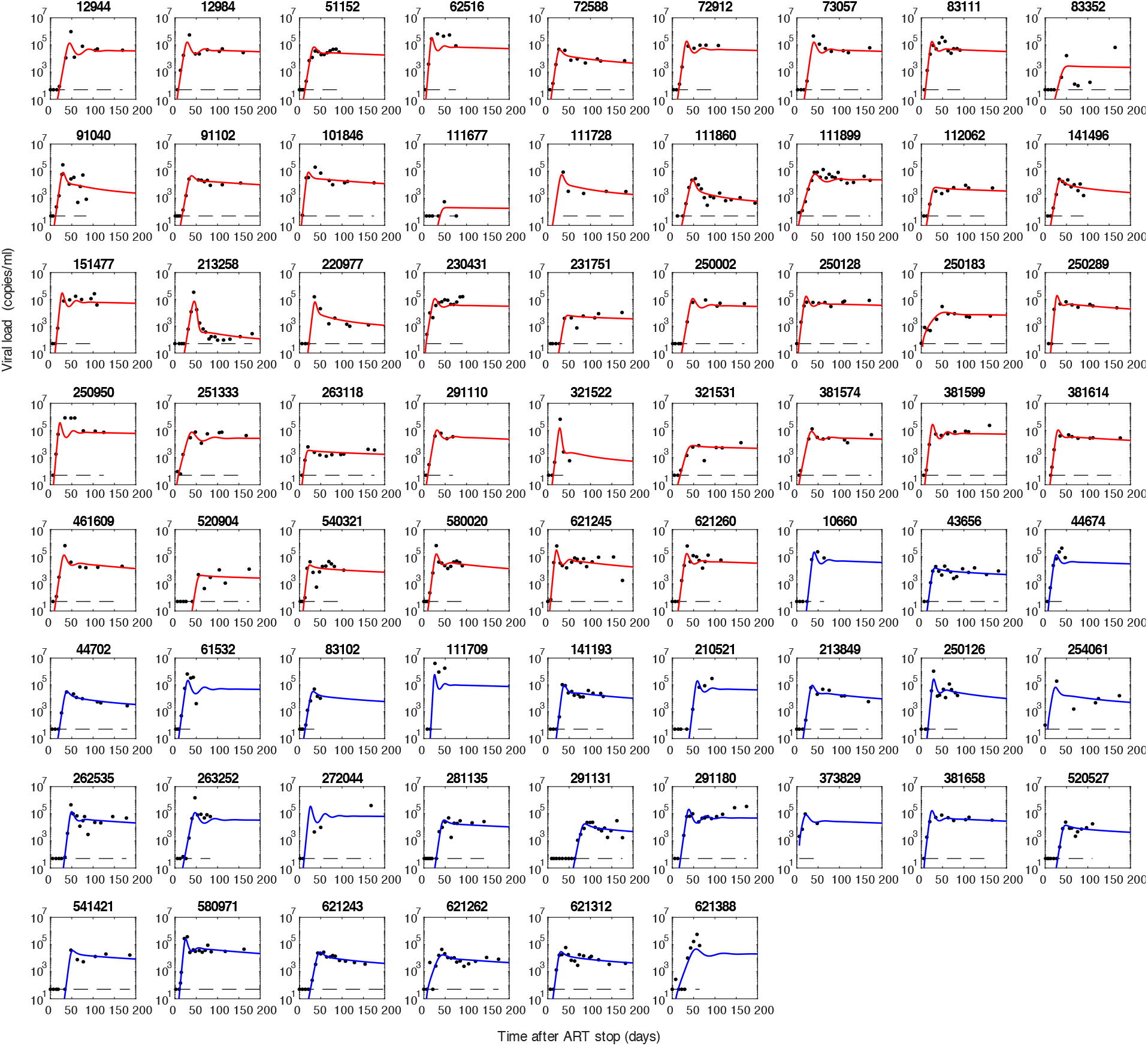
Viral load values and model fits during rebound for a cohort of HIV-infected individuals. Individuals in red were treated with non-NNRTI based antiretroviral therapy, while individuals in blue were treated with NNRTI-based regimes.

**Figure S8:**
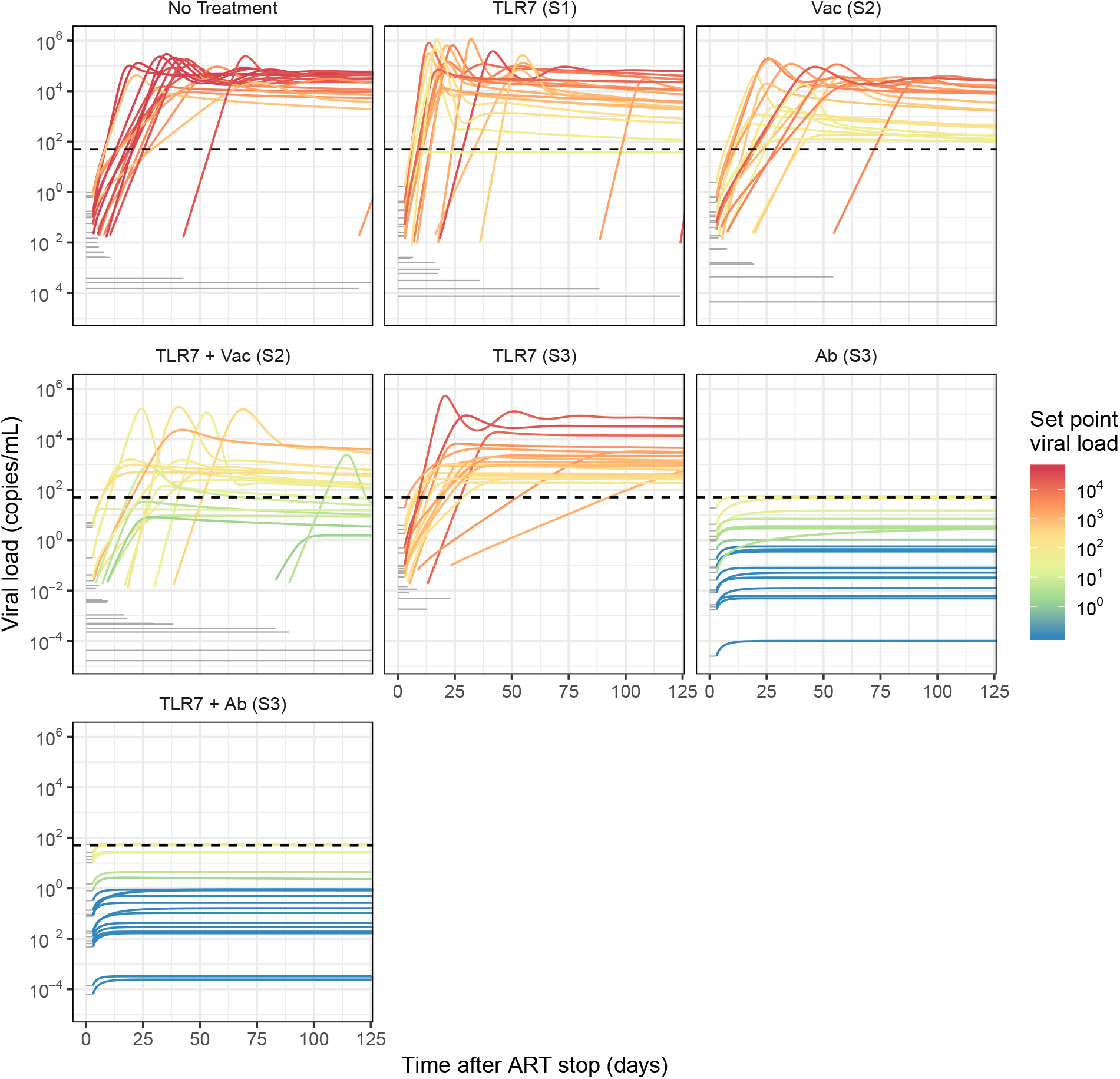
Simulated HIV rebound after hypothetical immunotherapeutic treatments in humans. In order to predict the effects of immunotherapies in multiple combinations in an HIV context, we simulated viral rebound trajectories by combining baseline population heterogeneity from human HIV rebound data with treatment effects from each study. To simulate an individual rebound trajectory, a set of parameters was sampled from the HIV population fitting results and then the relevant treatment effects were applied. The treatment effects applied in each panel correspond to the treatment effects identified in the indicated study; for clarity the list of effects can be found in Table S11. 20 example rebound trajectories are shown for each treatment. Viral load prior to succesful rebound is illustrated by a grey horizontal line at the level expected according to the simulated reservoir size. Each trajectory is colored according to its viral load at one year. Dotted lines indicate the detection threshold for standard viral load assays (50 copies/mL).

**Figure S9:**
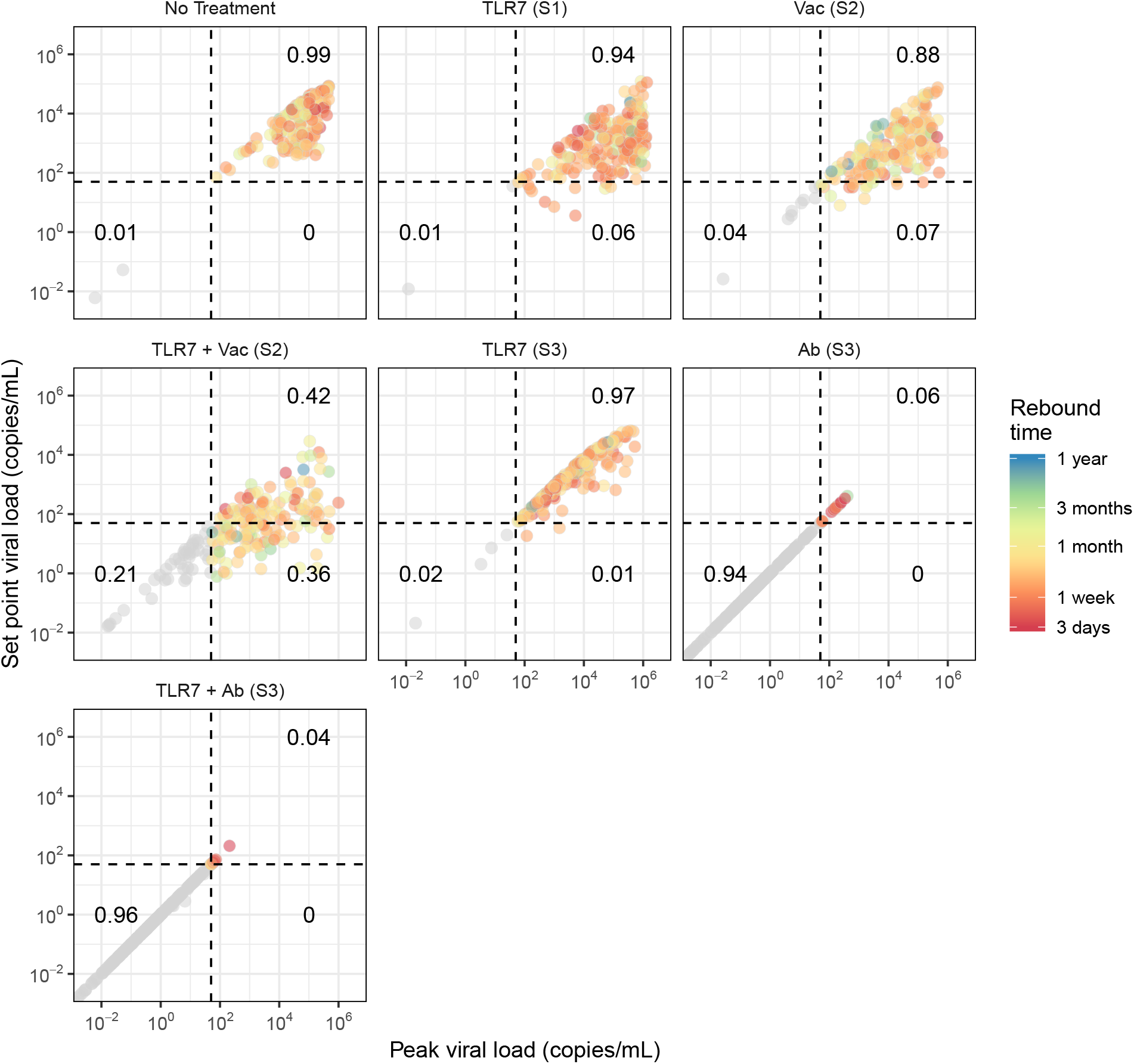
Summary statistics for simulated HIV rebound after alternative immunotherapeutic treatment in humans. 200 simulations for each treatment summarized by their peak and final viral loads. Each dot represents a particular individual and is colored according to the first time at which that individual rebounded (crossed 50 copies/mL). Individuals who never crossed this threshold are shown in gray. Dotted lines indicate the detection threshold for standard viral load assays (50 copies/mL) and dark black numbers show the proportion of individuals falling in the indicated quadrant. Viral rebound trajectories were simulated for one year.

**Figure S10:**
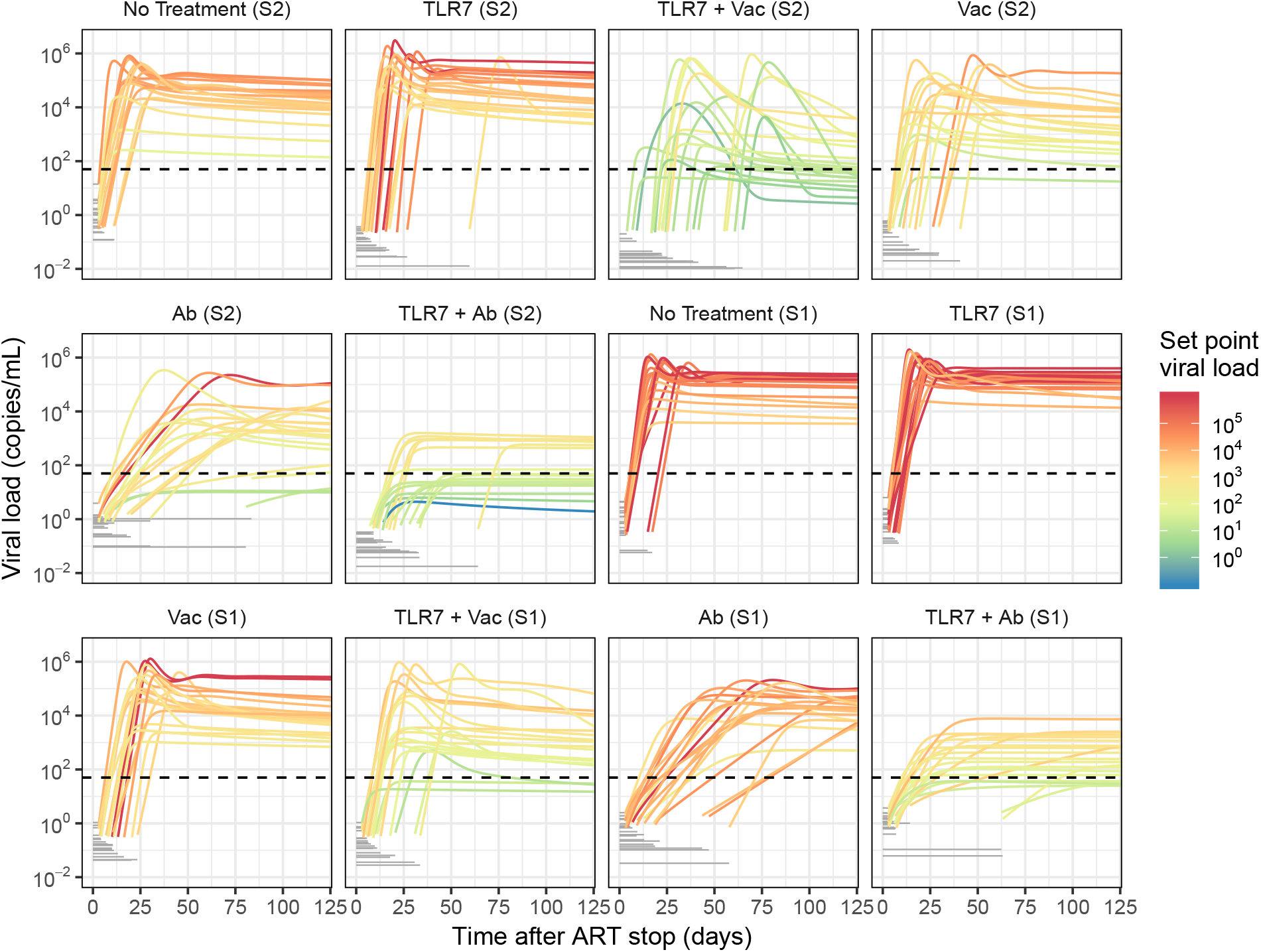
Simulated SIV rebound after alternative immunotherapeutic treatment in macaques. In order to predict the effects of immunotherapies in multiple combinations in the context of SIV, we simulated viral rebound trajectories by combining baseline population heterogeneity from macaque SIV rebound data (studies 1 and 2) with treatment effects from each study. To simulate an individual rebound trajectory, a set of parameters was sampled from the SIV population fitting results and then the relevant treatment effects were applied. The treatment effects applied in each panel correspond to the treatment effects identified in the indicated study; for clarity the list of effects can be found in Tables S12 and S13. 20 example rebound trajectories are shown for each treatment. Viral load prior to succesful rebound is illustrated by a grey horizontal line at the level expected according to the simulated reservoir size. Each trajectory is colored according to its viral load at one year. Dotted lines indicate the detection threshold for standard viral load assays (50 copies/mL).

**Figure S11:**
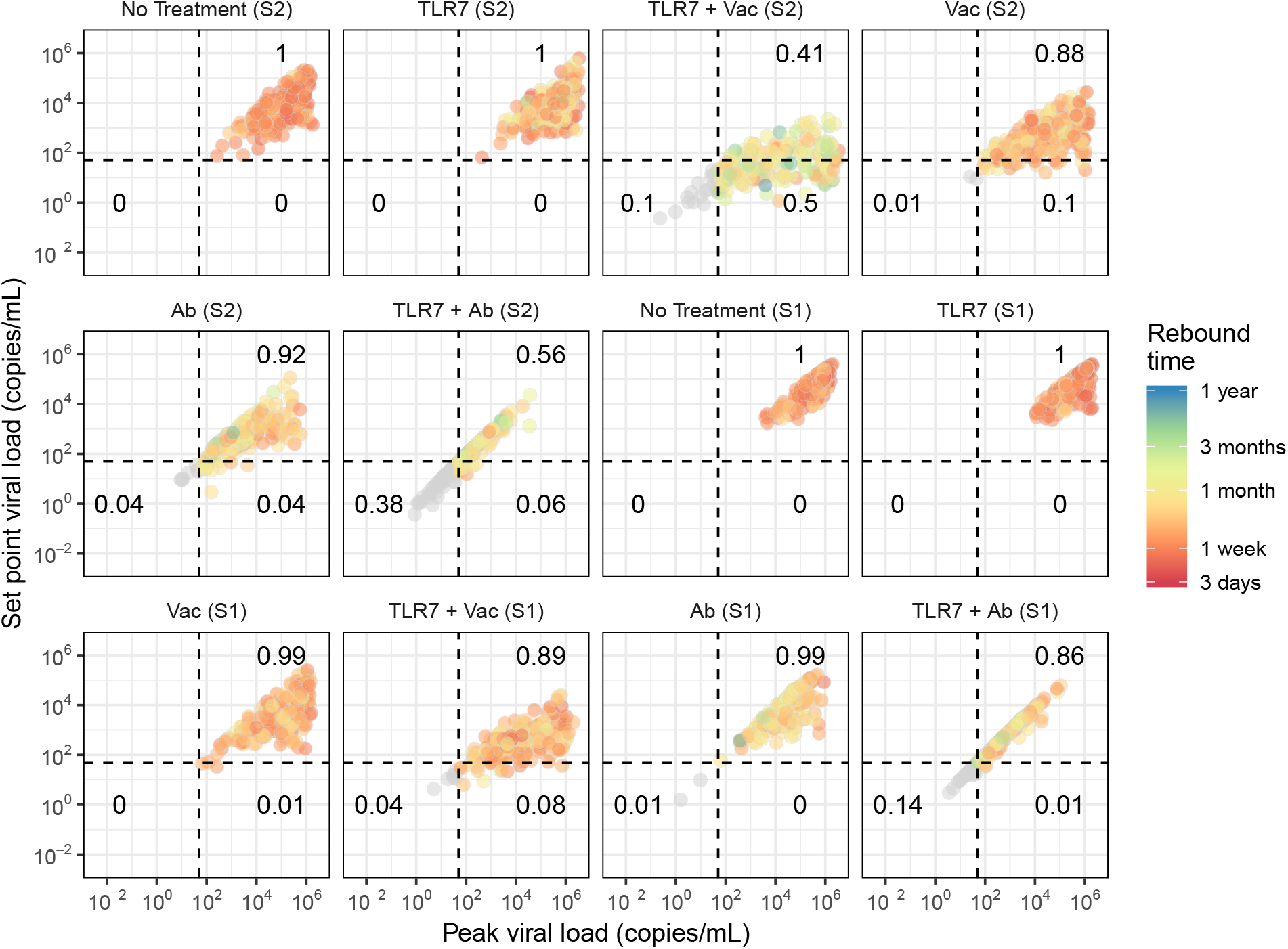
Summary statistics for simulated SIV rebound after alternative immunotherapeutic treatment in macaques. 200 simulations for each treatment summarized by their peak and final viral loads. Each dot represents a particular individual and is colored according to the first time at which that individual rebounded (crossed 50 copies/mL). Individuals who never crossed this threshold are shown in gray. Dotted lines indicate the detection threshold for standard viral load assays (50 copies/mL) and dark black numbers show the proportion of individuals falling in the indicated quadrant. Viral rebound trajectories were simulated for one year.

**Figure S12:**
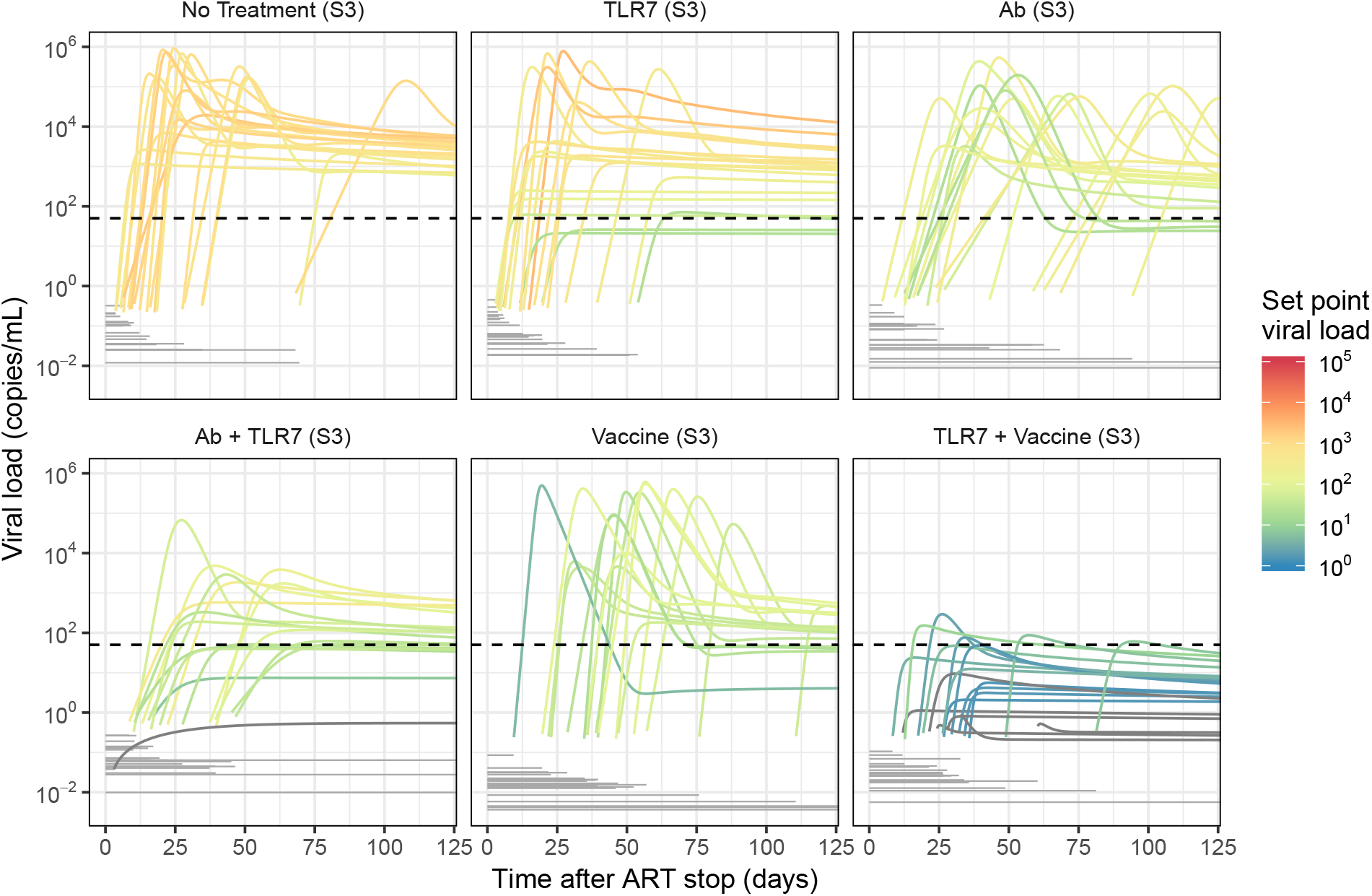
Simulated SHIV rebound after alternative immunotherapeutic treatment in macaques. In order to predict the effects of immunotherapies in multiple combinations in the context of SHIV, we simulated viral rebound trajectories by combining baseline population heterogeneity from macaque SHIV rebound data (study 3) with treatment effects from each study. To simulate an individual rebound trajectory, a set of parameters was sampled from the SHIV population fitting results and then the relevant treatment effects were applied. The treatment effects applied in each panel correspond to the treatment effects identified in the indicated study; for clarity the list of effects can be found in Table S14. 20 example rebound trajectories are shown for each treatment. Viral load prior to succesful rebound is illustrated by a grey horizontal line at the level expected according to the simulated reservoir size. Each trajectory is colored according to its viral load at one year. Dotted lines indicate the detection threshold for standard viral load assays (50 copies/mL).

**Figure S13:**
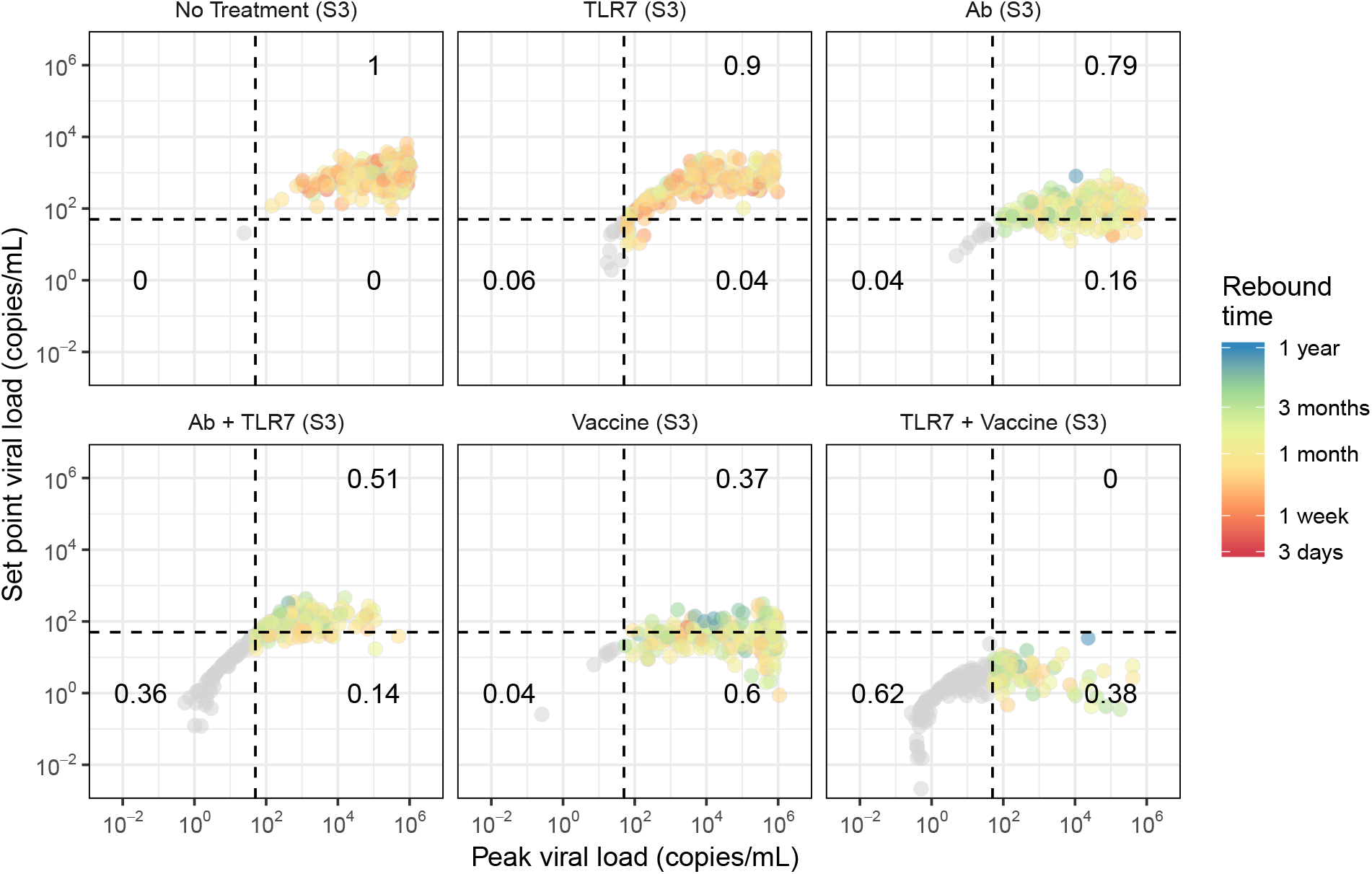
Summary statistics for simulated SHIV rebound after alternative immunotherapeutic treatment in macaques. 200 simulations for each treatment summarized by their peak and final viral loads. Each dot represents a particular individual and is colored according to the first time at which that individual rebounded (crossed 50 copies/mL). Individuals who never crossed this threshold are shown in gray. Dotted lines indicate the detection threshold for standard viral load assays (50 copies/mL) and dark black numbers show the proportion of individuals falling in the indicated quadrant. Viral rebound trajectories were simulated for one year.

**Table S3:**
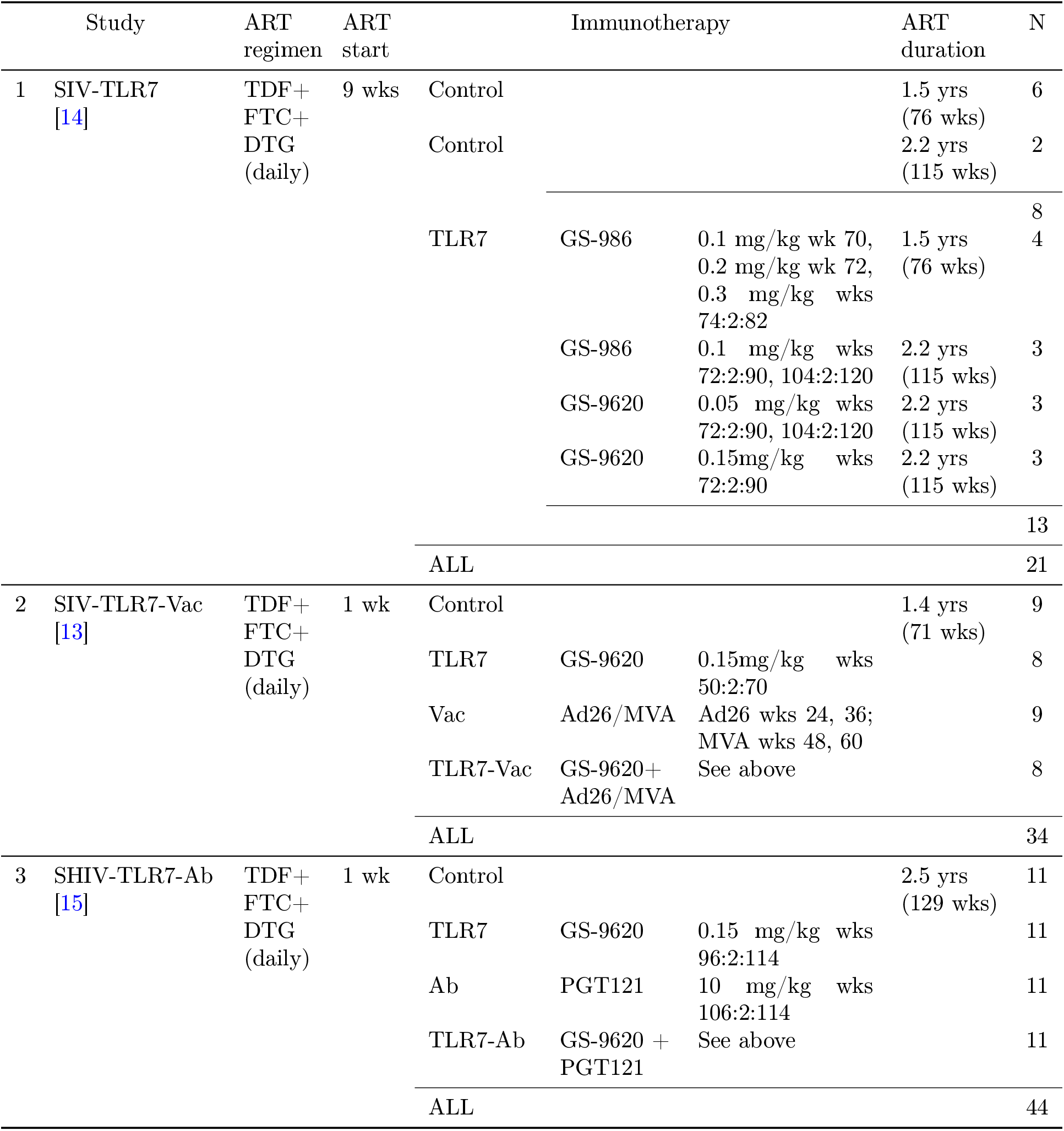
Summary of study designs.

**Table S4:**
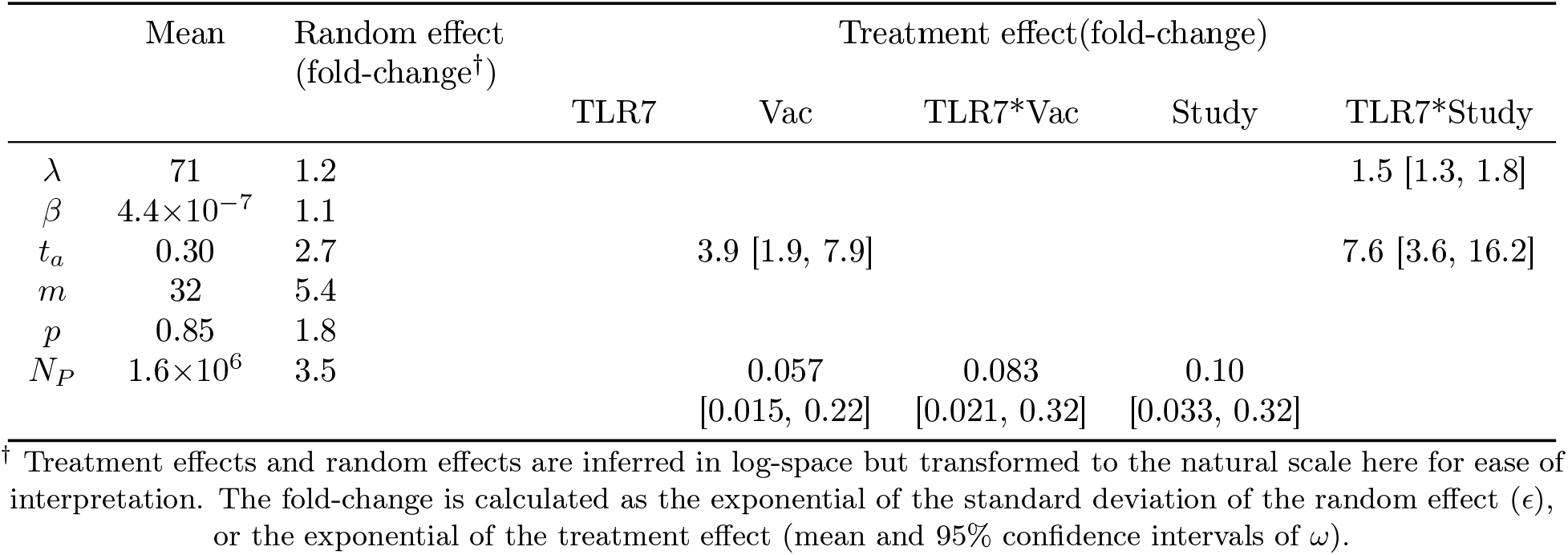
Population parameter values and treatment effects for best estimated model for SIV data (TRL7 ± Vac).

**Table S5:** Population parameter values and treatment effects for best estimated model for SHIV data (TRL7 ± Ab).

|  | Mean | Random effect<br>(fold-change<sup>†</sup>) | Treatment effect(fold-change) |  |  |
| --- | --- | --- | --- | --- | --- |
|  |  |  | TLR7 | Ab | TLR7*Ab |
| $\lambda$ | 39 | | | | |
| $\beta$ | $1.0 \times 10^{-6}$ | 1.3 | | 0.45 [0.34, 0.59] | |
| $t_a$ | 3.2 | 2.3 | | | |
| $m$ | 1.9 | 23 | 100 [6.1, 1700] | | |
| $p$ | 3.6 | | | | |
| $N_P$ | $7.7 \times 10^4$ | 1.9 | | 0.14 [0.061, 0.30] | |

<sup>†</sup> Treatment effects and random effects are inferred in log-space but transformed to the natural scale here for ease of interpretation. The fold-change is calculated as the exponential of the standard deviation of the random effect ( $\epsilon$ ), or the exponential of the treatment effect (mean and 95% confidence intervals of $\omega$ ).

**Table S6:**
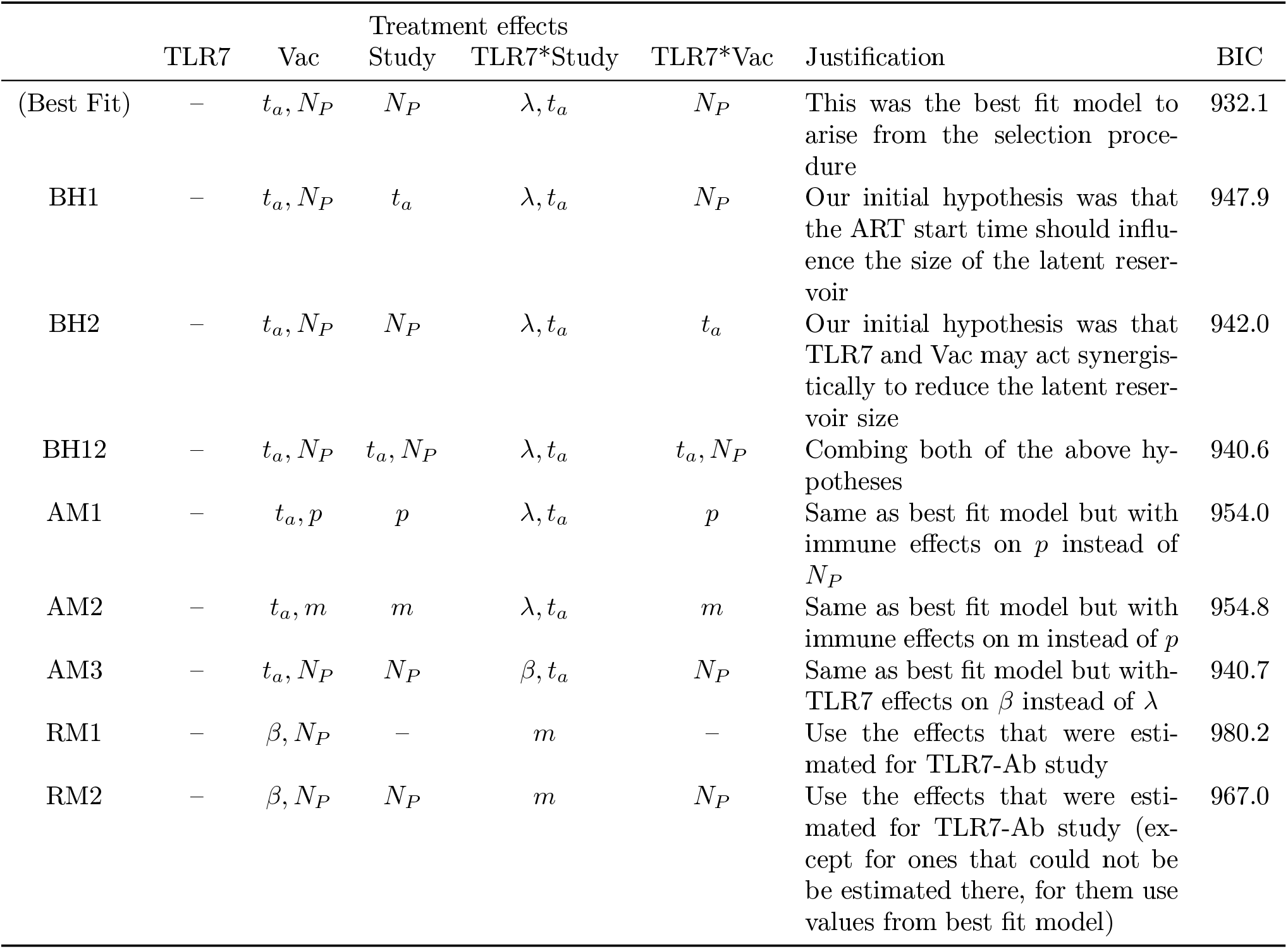
Alternative models tested for SIV data (TRL7 ± Vac). BIC = Bayesian Information Criterion

**Table S7:** Alternative models tested based for SHIV data (TRL7 ± Ab). BIC = Bayesian Information Criterion.

|  | Treatment effects |  |  | Justification | BIC |
| --- | --- | --- | --- | --- | --- |
|  | TLR7 | Ab | TLR7*Ab |  |  |
| (Best Fit) | $m$ | $\beta, N_P$ | – | This was the best fit model to arise from the selection procedure | 746.5 |
| BH1 | $m$ | $t_a, N_P$ | – | Our initial hypothesis was that the Ab would reduce the size of the latent reservoir | 763.7 |
| BH2 | $t_a$ | $\beta, N_P$ | – | Our initial hypothesis was that TLR7 would reduce the size of the latent reservoir, and, this effect was selected in the TLR7-Vac study | 766.0 |
| BH3 | $m$ | $\beta, N_P$ | $N_P$ | Our initial hypothesis is that the TLR7 may increase the effect of the Ab, and this interaction was observed for the vaccine in the TLR7-Vac study | 761.5 |
| AM1 | $N_P$ | $\beta, N_P$ | – | same as best fit model but including a TLR7 effect on $N_P$ instead of on $m$ | 760.7 |
| AM2 | $m$ | $\beta, m$ | – | same as best fit model but including a vaccine effect on $m$ instead of $N_P$ | 770.7 |
| AM3 | $\lambda$ | $\beta, N_P$ | – | same as best fit modeling but including a TLR7 effect on $\lambda$ instead of $m$, since $\lambda$ was selected for TLR7 in the TLR7-Vac study | 769.4 |
| AM4 | $m$ | $\lambda, N_P$ | – | same as best fit model but including a vaccine effect on $\lambda$ instead of on $\beta$ | 754.4 |
| RM1 | $\lambda, t_a$ | $t_a, N_P$ | – | Use the effects that were estimated for TLR7-Vac study | 747.5 |
| RM1b | $\lambda, t_a$ | $t_a, N_P$ | $t_a$ | Use the effects that were estimated for TLR7-Vac study plus $t_a$ interaction | 764.5 |
| RM2 | $\lambda, t_a$ | $t_a, N_P$ | $N_P$ | Use the effects that were estimated for TLR7-Vac study, including interaction | 778.3 |
| RM2b | $\lambda, t_a$ | $t_a, N_P$ | $t_a, N_P$ | Use the effects that were estimated for TLR7-Vac study, including interactions on both $t_a$ and $N_P$ | 776.6 |

**Table S8:** Details of TLR7-agnosist dosing variations received by animals in Study 2 [14].

| ID | ID # | ART duration<br>76 wks?<br>(vs 115 wks) | Description | GS-9620?<br>(vs GS-986) | # doses | Cumulative<br>dose<br>(mg/kg) | Average<br>dose<br>(mg/kg) | Max<br>dose<br>(mg/kg) |
| --- | --- | --- | --- | --- | --- | --- | --- | --- |
| 1 | 156-08 | 1 | GS-986 0.1 mg/kg | 0 | 7 | 1.8 | 0.26 | 0.3 |
| 2 | 166-08 | 1 | wk 70, 0.2 mg/kg | 0 | 7 | 1.8 | 0.26 | 0.3 |
| 3 | 280-09 | 1 | wk 72, 0.3 mg/kg | 0 | 7 | 1.8 | 0.26 | 0.3 |
| 4 | 310-09 | 1 | wks 74:2:82 | 0 | 7 | 1.8 | 0.26 | 0.3 |
| 5 | 205-08 | 1 | Control | 0 | 0 | 0 | 0 | 0 |
| 6 | 267-08 | 1 |  | 0 | 0 | 0 | 0 | 0 |
| 7 | 105-09 | 1 |  | 0 | 0 | 0 | 0 | 0 |
| 8 | 234-09 | 1 |  | 0 | 0 | 0 | 0 | 0 |
| 9 | 322-09 | 1 |  | 0 | 0 | 0 | 0 | 0 |
| 10 | 374-09 | 1 |  | 0 | 0 | 0 | 0 | 0 |
| 11 | 162-09 | 0 | Control | 0 | 19 | 0 | 0 | 0 |
| 12 | 305-10 | 0 |  | 0 | 19 | 0 | 0 | 0 |
| 13 | 280-10 | 0 | GS-986 0.1 mg/kg | 0 | 19 | 1.9 | 0.1 | 0.1 |
| 14 | 288-10 | 0 | wks 72:2:90, | 0 | 19 | 1.9 | 0.1 | 0.1 |
| 15 | 344-10 | 0 | 108:2:124 | 0 | 19 | 1.9 | 0.1 | 0.1 |
| 16 | 293-09 | 0 | GS-9620 0.05 | 1 | 19 | 0.95 | 0.05 | 0.05 |
| 17 | 295-10 | 0 | mg/kg wks 72:2:90, | 1 | 19 | 0.95 | 0.05 | 0.05 |
| 18 | 304-10 | 0 | 108:2:124 | 1 | 19 | 0.95 | 0.05 | 0.05 |
| 19 | 177-10 | 0 | GS-9620 | 1 | 10 | 1.5 | 0.15 | 0.15 |
| 20 | 341-10 | 0 | 0.15mg/kg wks | 1 | 10 | 1.5 | 0.15 | 0.15 |
| 21 | 412-10 | 0 | 72:2:90 | 1 | 10 | 1.5 | 0.15 | 0.15 |

**Table S9:** Comparing population parameter values in the absense of treatment for SIV vs SHIV vs HIV. Parameter meanings and units are given in the **SI Methods**. Parameters not listed are assumed to be the same between viruses and are given in Tables S1 and S2.

| Parameter | SIV |  | SHIV |  | HIV |  |
| --- | --- | --- | --- | --- | --- | --- |
|  | Mean | Random effect<br>(fold-change) | Mean | Random effect<br>(fold-change) | Mean | Random effect<br>(fold-change) |
| **Fit:** |  |  |  |  |  |  |
| $\lambda$ | 71 | 1.2 | 39 | | 330 | 1.2 |
| $\beta$ | $4.4 \times 10^{-7}$ | 1.1 | $1.0 \times 10^{-6}$ | 1.3 | $5.30 \times 10^{-7}$ | 1.1 |
| $t_a$ | 0.30 | 2.7 | 3.2 | 2.3 | 0.014 | 20 |
| $m$ | 32 | 5.4 | 1.9 | 23 | 3.8 | 12 |
| $p$ | 0.85 | 1.8 | 3.6 | | 3.3 | 2.7 |
| $N_P$ | $1.6 \times 10^6$ | 3.5 | $7.7 \times 10^4$ | 1.9 | $9.3 \times 10^5$ | 4.4 |
| **Fixed:** |  |  |  |  |  |  |
| $d_I$ | 0.4 | | 0.4 | | 1 | |
| $k$ | 50,000 | | 50,000 | | 10,000 | |
| $I_1$ | $4 \times 10^{-5}$ | | $4 \times 10^{-5}$ | | $4 \times 10^{-6}$ | |

**Table S10:** Human main figure simulated treatment effects (Figure 5).

| Panel name | Treatment effects |
| --- | --- |
| No Treatment | None |
| TLR7 (S1) | TLR7xStudy:$\lambda$, TLR7xStudy:$t_a$ |
| TLR7 + Vaccine (S2) | Vac:$t_a$, Vac:$N_P$, TLR7xVac:$N_P$ |
| TLR7 + Antibody (S3) | Ab:$\beta$, Ab:$N_P$, TLR7:$m$ |

**Table S11:** Human supplemental figure simulated treatment effects (Figures S8, S9).

| Panel Name | Treatment effects included |
| --- | --- |
| No Treatment | None |
| TLR7 (S1) | TLR7xStudy:$\lambda$, TLR7xStudy:$t_a$ |
| Vaccine (S2) | Vac:$t_a$, Vac:$N_P$ |
| TLR7 + Vaccine (S2) | Vac:$t_a$, Vac:$N_P$, TLR7xVac:$N_P$ |
| TLR7(S3) | TLR7:$m$ |
| Ab (S3) | Ab:$\beta$, Ab:$N_P$ |
| TLR7 + Antibody (S3) | Ab:$\beta$, Ab:$N_P$, TLR7:$m$ |

**Table S12:** SIV Study 1 simulated treatment effects (Figures S10, S11).

| Panel Name | Treatment effects included |
| --- | --- |
| No treatment (S1) | Study:$N_P$ |
| TLR7 (S1) | Study:$N_P$, TLR7xStudy:$\lambda$, TLR7xStudy:$t_a$ |
| TLR7 + Vac (S1) | Study:$N_P$, TLR7xStudy:$\lambda$, TLR7xStudy:$t_a$, Vac:$t_a$, Vaccine:$N_P$, TLR7xVac:$N_P$ |
| Vac (S1) | Study:$N_P$, Vac:$t_a$, Vac:$N_P$ |
| Ab (S1) | Study:$N_P$, Ab:$\beta$, Ab:$N_P$ |
| TLR7 + Ab (S1) | Study:$N_P$, Ab:$\beta$, Ab:$N_P$, TLR7xStudy:$t_a$, TLR7xStudy:$\lambda$ |

**Table S13:** SIV Study 2 simulated treatment effects (Figures S10, S11).

| Panel Name | Treatment effects included |
| --- | --- |
| No treatment (S2) | None |
| TLR7 (S2) | None |
| Vac (S2) | Vac:$t_a$, Vac:$N_P$ |
| TLR7 + Vac (S2) | Vac:$t_a$, Vac:$N_P$, TLR7xVac:$N_P$ |
| Ab (S2) | Ab:$\beta$, Ab:$N_P$ |
| TLR7 + Ab (S2) | Ab:$\beta$, Ab:$N_P$ |

**Table S14:** SHIV simulated treatment effects (Figures S12, S13).

| Panel Name | Treatment effects included |
| --- | --- |
| No treatment (S3) | None |
| TLR7 (S3) | TLR7:$m$ |
| Ab (S3) | Ab:$\beta$, Ab:$N_P$ |
| TLR7 + AB (S3) | TLR7:$m$, Ab:$\beta$, Ab:$N_P$ |
| Vac (SS) | Vac:$t_a$, Vac:$N_P$ |
| Vac + TLR7 (S3) | Vac:$t_a$, Vac:$N_P$, TLR7:$m$ |

